# An aphid effector pair forms a hetero-oligomeric complex important for protein stability and activity

**DOI:** 10.64898/2026.09.18.752295

**Authors:** Thomas Waksman, Jade R. Bleau, Ronan S. Fisher, Genady Pankovs, William N. Hunter, Jorunn I.B. Bos

## Abstract

During disease and infestation, pathogens and pests deliver effectors inside their host to manipulate host responses, leading to immune suppression and altered nutrient availability. To be successful, pathogens and pests can produce and secrete large repertoires of effectors, consisting of up to hundreds of proteins. While effector virulence activities are usually studied in isolation, effectors can function together or cooperate to regulate transport and/or activity. An important example is the suppression of effector avirulence activity by plant pathogenic microbe effectors. Moreover, some effectors can physically associate with target host proteins and promote virulence. Here we explored the interaction of a conserved and co-regulated effector pair in aphids. Using a combination of computational modelling and crystallography combined with functional assays, we reveal a novel type of oligomeric assembly of the paired aphid effectors in a hexameric/octameric complex. Our data indicate that different oligomerization states exist among orthologs of this effector complex, suggesting that dynamic dissociation or subunit exchange may occur. Structure-based mutagenesis revealed that complex formation is essential for both effector protein stability and the *in planta* activity of one of these proteins. These findings emphasize the significance of effector hetero-oligomerization at the plant-insect interface, consistent with the model that this effector pair is an evolutionarily conserved module across aphid species.

## Introduction

Pathogens and pests deliver effector molecules, including RNA and proteins, inside their host to suppress host immune responses and facilitate access to nutrients for life cycle support leading to disease and infestation. Once delivered, these effectors interact with host proteins as well as other molecules and modify their activity as part of an effective virulence strategy (1). However, as a protective mechanism these effectors may be recognised by immune receptors, either through direct protein-protein interactions, or indirectly through detection of effector activity, leading to activation of host immunity and resistance (2). Unveiling the complexity of host and pathogen or pest interkingdom interactions strengthens our understanding of the molecular dialogues that determine disease and underpins development of disease and infestation prevention and control strategies.

While effector protein characterization typically focuses on one protein at a time, effectors may feature metaeffector activity, whereby effectors modulate the activity of other effectors either indirectly, by targeting similar host target proteins or pathways, or directly through physical protein-protein interaction. For example, LubX from *Legionella pneumophila*, a ubiquitin E3 ligase, targets another effector, SidH, for proteasome-mediated degradation to regulate its activity during infection stages (3). Moreover, ≈20 interacting protein pairs were identified through interaction analyses of ≈330-390 *L. pneumophila* effectors pointing to extensive metaeffector activity (4, 5). Similarly, effectors from plant pathogenic microbes can feature metaeffector activity, with an evident example being the suppression of Effector-Triggered-Immunity, where one effector suppresses the immune receptor-mediated activation of another effector (6). Also, effectors from plant pathogenic microbes can form heteromeric complexes, and although their biological function remains to be elucidated, these are suggested to be involved in either regulation of effector translocation and/or activity (7–11). For instance, the effector Six5 from *Fusarium oxysporum f. sp. lycopersici* (Fol) interacts with Avr2 at the plasmodesmata (PD), thereby increasing the PD size exclusion limit. This interaction facilitates the symplastic transport of Avr2 and potentially other effectors (8, 12). Five effectors from the fungus *Ustilago maydis* form a cell surface-exposed complex with two fungal membrane proteins, which is required for host colonization (10). Many pathogen and pest effector repertoires feature an over-representation of unknown and disordered proteins, and complex formation may be an important and widespread mechanism to diversify and regulate effector activities that determine virulence strategies (13–15).

We recently discovered that two aphid effectors encoded by a highly conserved, genetically linked and co-expressed gene pair form a complex that targets a host cell trafficking protein (16). Although these effectors physically interact, functional characterization of the individual proteins from the economically important aphid pests *Myzus persicae* and *Rhopalosiphum padi* pointed to important virulence activities (17, 18). Specifically, Mp1, from *M. persicae*, can associate with Vacuolar Protein Sorting associated Protein 52 (VPS52) in host but not nonhost plants (17). StVPS52 ectopic expression in the host plant *Nicotiana benthamiana* reduced susceptibility to aphids, and host plant infestation reduced detectable VPS52 protein levels, indicating that VPS52 is an important virulence target. Ectopic expression of both Mp1 and a putative ortholog from *R. padi*, Rp1, in host plants enhanced host susceptibility, pointing to an important role of this effector in aphid infestation (17–19). In contrast, ectopic expression of the other member of the effector pair, Mp58 from *M. persicae*, in host plants reduced susceptibility to *M. persicae*, suggesting this effector may trigger plant defence responses (18, 20). However, these phenotypes observed from ectopic expression can vary depending on the expression system used. In contrast, the Mp58-like effector from *Macrosiphum euphorbiae*, called Me10, promotes plant susceptibility (21) and interacts with a tomato 14-3-3 isoform 7 (TFT7), which contributes plant immunity (22).

Here, we explored the co-evolution and structural properties of the conserved and co-regulated aphid effector pair, based on Mp1-Mp58 and Rp1-Rp58, and the complex these effectors form. We used a phylogenetic analysis, structural biology, computational modelling, as well as mutagenesis and *in planta* assays to show that the aphid effector pair form a hetero-octameric complex. The 3D effector complex structure, in which Mp58/Rp58 forms a core tetramer, showed no similarity to structures in any protein structure database, suggesting that the complex is novel. Complex formation takes place in aphid saliva, upon ectopic expression in plants, as well as when proteins are produced *in vitro*. Mutations that impair the formation of the effector complex also led to reduced Mp1 and Mp58 stability, along with a reduction in Mp58 effector activity. Our findings show that effector hetero-oligomerization at the plant–insect interface is important, consistent with this effector pair functioning as an evolutionarily conserved module across aphid species.

## Results

### The Mp1-Mp58 effector module is highly conserved in aphids

To determine whether Mp1- and Mp58-like sequences potentially co-evolved, we performed phylogenetic analyses of homologues across 28 aphid species. Sequence similarity searches only identified Mp1- and Mp58-like sequences within the *Aphididae*, but not in any other Hemipteran species. In both trees, terminal relationships were generally better supported than deeper nodes indicating conservation among closely related homologues but uncertainty between more divergent lineages (**Figure 1**). In both the Mp1 and Mp58 phylogenies, the *M. persicae* and *Myzus varians* (equences formed a sister pair within a grouping that also contained the strongly supported *Brachycaudus klugkisti* and *Brachycaudus helichrysi* sequences (**Figure 1**). Visual TreeCmp comparison of the Mp1 and Mp58 maximum-likelihood trees showed substantial topological similarity across 28 shared taxa. The trees differed by a Robinson-Foulds distance of 10, which corresponds to a normalized RF distance of 0.20. Of 20,475 quartets 2,084 were discordant, with a normalized quartet distance of 0.102, indicating that 89.8% of four-taxon relationships were conserved. This supports broadly parallel phylogenetic histories and co-conservation of Mp1 and Mp58 as a putative paired effector module. Pairwise patristic distance matrices from Mp1 and Mp58 trees were compared using a Mantel test with Spearman correlation, accounting for the non-independence of pairwise distances. Detected evolutionary distances were strongly positively correlated Mantel r = 0.907, P = ≤ 0.0001; 9,999 permutations which indicates that taxa which separated by greater distances in the Mp1 phylogeny also tended to be more divergent in the Mp58 phylogeny. Although these observations support molecular co-evolution of the effector pair, they may also partly reflect the underlying aphid species phylogeny.

**Figure 1.**
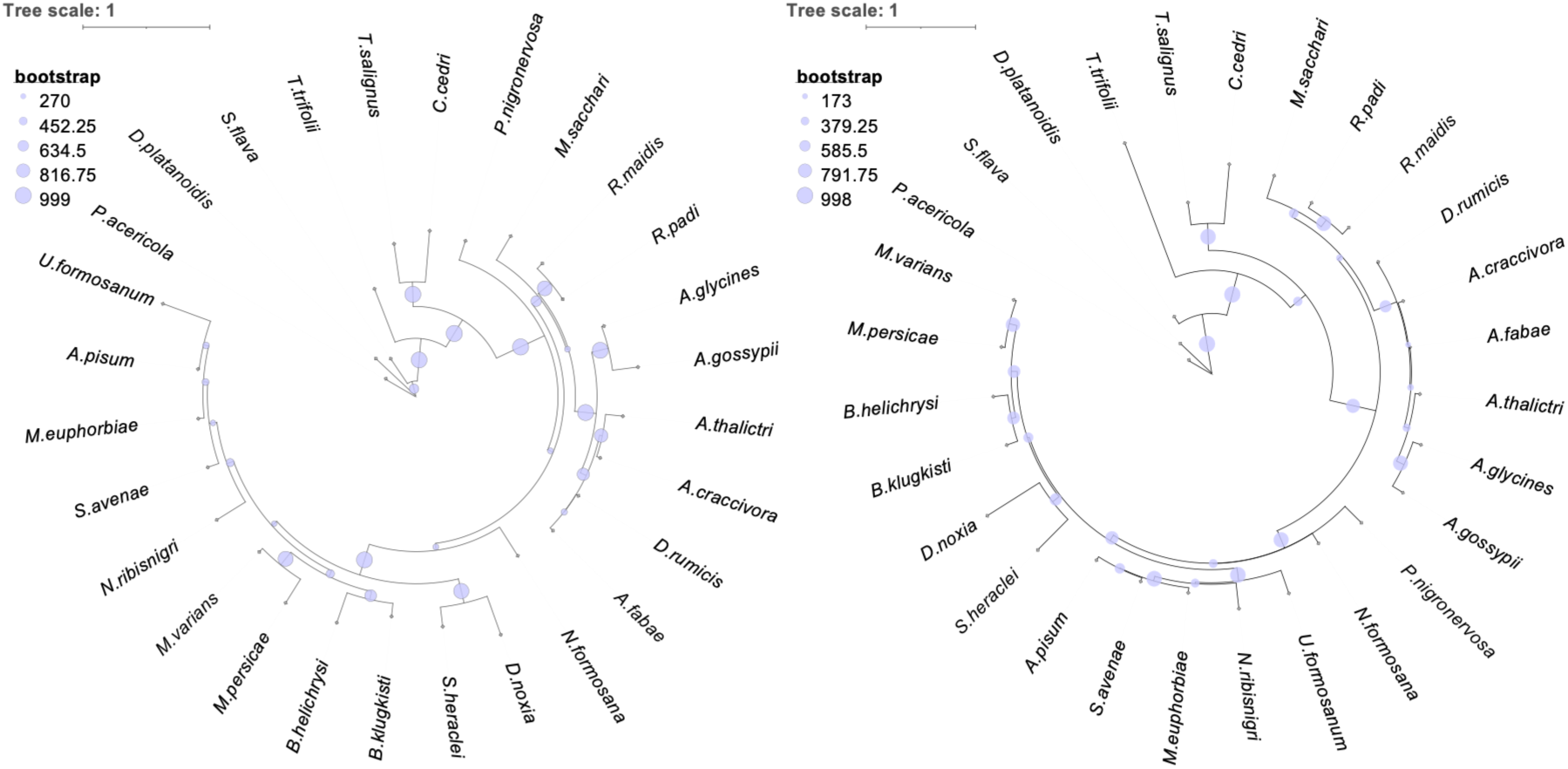
Phylogenetic trees generated for the Mp1- and Mp58-like family in 28 aphid species. Maximum-likelihood phylogenetic tree diagrams for Mp1 (left) and Mp58 (right). Lilac circles at internal nodes indicate their normalized transfer bootstrap expectation support values, with diameter proportional to support. Branch lengths are proportional to substitutions per site.

### Effector Mp58 forms a tetrameric structure

To determine the physical properties and structure of Mp58, we produced recombinant protein for Mp58, and the putative ortholog Rp58, in *E. coli*. While the theoretical mass of Mp58 and Rp58 is ≈15 kDa, size exclusion chromatography with multi-angle light scattering (SEC-MALS) measured a mass of ≈60 kDa (**Figure 2A**, **supplementary information**), suggesting that these proteins form a tetrameric assembly. Digestion with a low-specificity protease (chymotrypsin) produced a tetrameric species of lower mass (≈45 kDa), consistent with the core tetramer (*i.e.* full-length protein minus the protruding C-terminal region) in the AlphaFold structure prediction (**Figure 2B**, **supplementary information**). Chymotrypsin-digested Rp58 readily crystallized (**Figure S1A**) and diffraction data extending to 1.7 Å resolution were obtained. A Matthews coefficient of 2.39 Å³ Da⁻¹ supported a homotetramer in the asymmetric unit with a solvent content of around 50 % by volume. The crystal structure analysis was initiated using the AlphaFold prediction of Rp58_3-83_ (*i.e.* core tetramer, C-terminal region removed) for phasing by molecular replacement.

**Figure 2.**
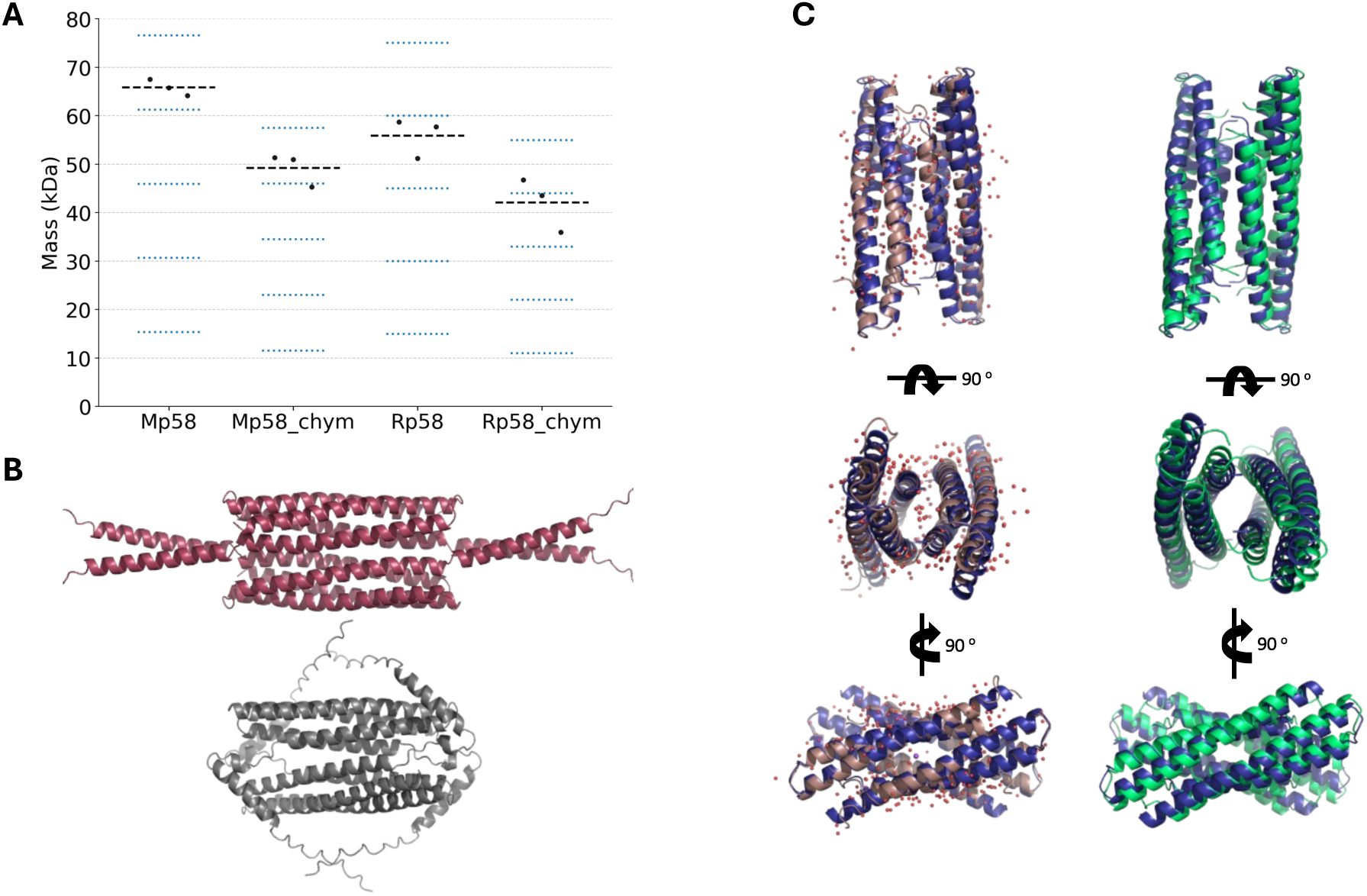
Structure of aphid effectors Mp58/Rp58. **(A)** Size exclusion chromatography with multi-angle light scattering (SEC-MALS) results for Mp58 and Rp58 proteins, expressed in and purified from *E. coli*, digested with tobacco etch virus protease (no suffix) or chymotrypsin (“_chym” suffix). Graph shows the mass measured for 3 biological replicates (black dots), mean mass (black dashed line), expected mass of oligomerization states 1-5 (blue dashed lines). **(B)** AlphaFold 3 models, shown in PyMOL cartoon style for tetrameric full-length Mp58 (red) and Rp58 (grey). **(C)** protein models superimposed using PyMOL “super” command for the Rp58 chymotrypsin fragment crystal structure (brown, with red dots for waters), molecular replacement search model (tetrameric Rp58_3-83_ AlphaFold 3 prediction in blue), tetrameric Mp58_3-83_ AlphaFold 3 prediction (green). Models are shown in PyMOL cartoon style in three orientations rotated by 90°.

Several rounds of electron and difference density map inspections, model building and fitting, location of solvent molecules interspersed with least-squares refinement, combined with using PDB-redo (23) resulted in a model (**Figure 2C, S1B**). The R factors (R_work_ = 0.21, R_free_ = 0.25) are lower than average for this resolution, reflecting the presence of heterogeneous termini from protease digestion, anisotropic diffraction and translational non-crystallographic symmetry. The AlphaFold model used for molecular replacement has a high confidence score and is similar to the crystal structure, with the only differences being in amino acid side chain conformation (AlphaFold pTM = 0.88, TM score = 0.95) (**Figure 2C**). Moreover, the AlphaFold prediction of the Mp58 core has a very similar structure to that of the Rp58 core (TM score = 0.90) (**Figure 2C**, **S2**). Overall, these data indicate that the structures of Mp58 and Rp58 (experimental or predicted) are accurate to reasonably high resolution.

### Effector oligomerization in saliva and plants

Since we found that Mp58 forms a homo-tetramer *in vitro*, we wanted to determine whether Mp58 is also present in an oligomeric state upon secretion into aphid saliva. We collected saliva from *M. persicae* and performed SDS-PAGE and BN-PAGE (Blue Native PAGE) followed by immunoblotting with a native Mp58 antibody. Upon SDS-PAGE, we detected Mp58 at its expected size (≈15 kDa) in saliva, alongside purified Mp58 (**Figure 3A**). However, upon BN-PAGE we detected several slower migrating bands at and above ≈146 kDa (**Figure 3A**) suggesting that Mp58 exists in a high molecular weight complex upon secretion, either in homo- and/or hetero-oligomeric protein complexes with other salivary proteins. Mp58 purified from *E. coli* migrated between the 66 and 146 kDa markers upon BN-PAGE and immunoblotting, consistent with its SEC mobility and tetrameric state (**Figure 3B**).

**Figure 3.**
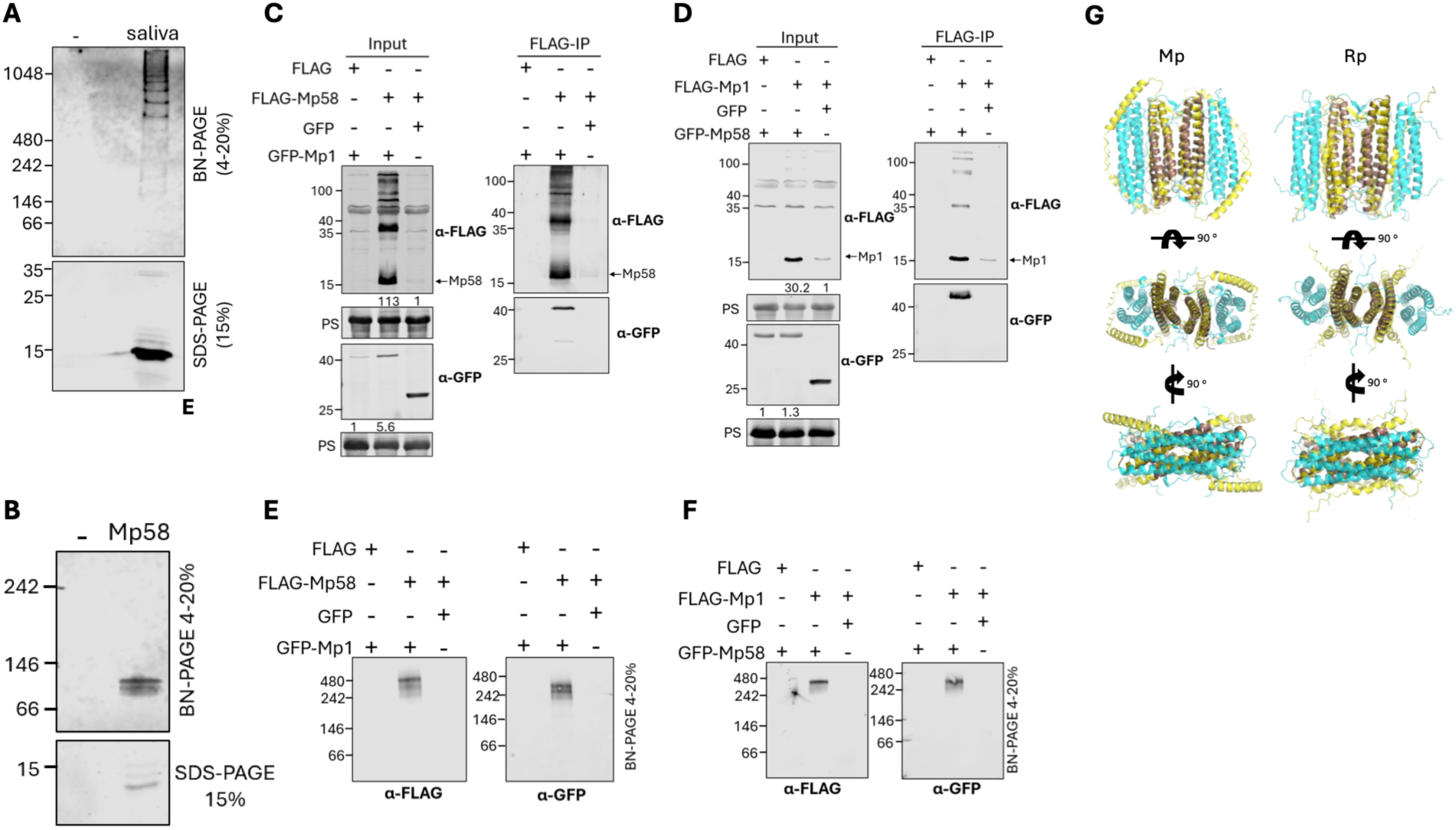
Oligomeric Mp1-Mp58 effector complex formation *in vivo* and predicted its structure. **(A-F)** Immunoblotting (IB) after polyacrylamide gel electrophoresis (PAGE) using either sodium dodecyl sulfate (SDS) or blue native (BN) conditions. Numbers refer to mass of PAGE marker proteins (kilodaltons, left of image) or protein quantification (below image). **A,** Mp58 IB using *Myzus persicae* saliva or aphid-free saliva collection solution (-). **B,** Mp58 IB using Mp58 expressed in and purified from *E. coli* or buffer (-). **(C-F)** IB of Mp58 and Mp1 expressed with either flag or green fluorescent protein (GFP) tags in *Nicotiana benthamiana*. SDS-PAGE was used before (input) or after immunoprecipitation with FLAG-beads (FLAG-IP) (**C, D**); BN-PAGE was used after FLAG-IP to detect native protein complexes (**E, F**). Protein quantification in panels **C** C **D** was performed by normalising the band intensity of FLAG-Mp1/Mp58 and GFP-Mp1/Mp58 to the total protein amounts using Empiria Studio and LI-COR images. FLAG-Mp1/Mp58 levels in samples co-expressed with GFP-Mp58/Mp1 were compared to protein levels in samples co-expressed with GFP (control, set at 1) to generate band intensity ratios. GFP-Mp1/Mp58 levels in samples co-expressed with FLAG-Mp58/Mp1 were compared to protein levels in samples co-expressed with FLAG (control, set at 1) to generate band intensity ratios. PS indicates Ponceau Stained blots showing loading and transfer. **(G)** protein models superimposed using PyMOL “super” command for the Rp58 chymotrypsin fragment crystal structure (brown), and octameric AlphaFold 3 prediction of either the *Myzus persicae* (Mp) or *Rhopalosiphum padi* (Rp) effector complex with 4 Mp58/Rp58 (yellow) and 4 Mp1/Rp1 (cyan) subunits. Models are shown in PyMOL cartoon style in three orientations rotated by 90°. Data are representative of n = 2 independent biological replicates.

As we previously found that Mp58 and Mp1 physically interact (16), we investigated the oligomeric state of Mp58 and the Mp1-Mp58 complex upon co-expression *in planta*. We transiently co-expressed FLAG-Mp58 with GFP-Mp1 or FLAG-Mp1 with GFP-Mp58 in *N. benthamiana* and performed co-immunoprecipitation. We detected a faint band corresponding to FLAG-Mp58 or FLAG-Mp1 based on their expected monomeric size (≈15kDa) when these proteins were expressed with the GFP control (**Figure 3C, 3D**). Both proteins showed an increase in abundance when co-expressed with GFP-Mp1 or GFP-Mp58, respectively. We also observed an increase in GFP-Mp1 abundance in the presence of FLAG-Mp58 (**Figure 3C, S3A**). In line with our previous observations, Mp58 and Mp1 co-immunoprecipitated indicative of the formation of an Mp1-Mp58 effector complex. BN-PAGE followed by immunoblotting using the same samples showed a double band corresponding to the effector complex migrating between the 242 kDa and 480 kDa markers (**Figure 3E, 3F**). We were unable to detect FLAG-Mp1 and FLAG-Mp58 by BN-PAGE when expressed alone, which is likely due to low protein abundance of these individual effectors (**Figure 3E, 3F**). Nevertheless, the band corresponding to the Mp1-Mp58 complex was slower migrating than GFP-Mp1 or GFP-Mp58 alone (**Figure S3B**), pointing to the formation of a larger hetero-oligomeric Mp1-Mp58 complex in plants.

### Mp1 and Mp58 are present in an octameric effector complex

With our data supporting the formation of a large hetero-oligomeric Mp1-Mp58 effector complex, we were interested in investigating the 3D protein structure and complex stoichiometry. Several observations suggested a stoichiometry of 4+4 for the Mp1-Mp58 complex and Rp1-Rp58, although other oligomerization states or conformations may occur. The BN-PAGE data supports the effectors associating in a 1:1 molar ratio (the effectors share a similar size of 15 kDa), while both SEC-MALS and crystallographic analysis support a tetrameric state for Mp58/Rp58. To determine stoichiometry experimentally, we used SEC-MALS to measure the mass of effector complexes formed by effectors with different tags. Despite substantial variability, which may be due to inherent dynamics of the effector complex, the results supported that a 4+4 or 4+2 stoichiometry is likely to predominate. For Rp58+Rp1, the mass with or without maltose binding protein (MBP) tag on Rp1 was close to the theoretical mass of a 4+2 complex (**Figure S4**, **supplementary information**). For Mp58+Mp1, the data was more variable but nonetheless the data points were close to the theoretical mass of a 4+2 or 4+4 complex (**Figure S4**, **supplementary information**). These two stoichiometries (4+2 and 4+4) may correspond to the double band observed in BN-PAGE (**Figures 3E, 3F**). Considering the combined SEC, BN-PAGE and structural data, an asymmetric complex involving an odd number of monomers of either protein is less likely.

Based on the experimentally determined stoichiometry and supported by the crystal structure data for Rp58, we generated AlphaFold predictions of the effector complex structures using a 4+4 stoichiometry, which is equivalent to the 4+2 structure with 2 extra Mp1 or Rp1 monomers that provide symmetry for visual clarity (**Figure 3G**). The structure of both orthologous effector complexes consists of two Mp58/Rp58 dimers at the core of the complex, in the same conformation as in the crystal structure, plus 2 Mp1/Rp1 dimers on either side (**Figure 3G**, **S2**). Each dimer is made of two antiparallel monomers, each monomer consisting of two antiparallel α-helices; the angle between the Mp58/Rp58 dimer and the Mp1/Rp1 dimer is lower than the angle between the two Mp58 or Rp58 dimers (**Figure 3G**). The subtle conformational differences between sub-complexes within the octamer, as well as differences between homotetramer structures within or without the octamer context are reflected in TM score when the models are aligned computationally (**Figure S2**). Amino acid conservation is lowest in the dimer-dimer interface within Mp58 or Rp58, being higher for all other dimer-dimer or monomer-monomer interfaces within the protein (**Figure S5**, **supplementary information**). AlphaFold confidence is higher for the heterotetramer half of the octamer than the homotetramer core (*i.e.* 3 points higher average pLDDT of ɑ-helix regions) for both effector complex orthologues (**Table S1**). The described oligomeric fold appears to be novel, as no matches were found using Foldseek Multimer in searches for similar existing structures, including sub-structures such as the Mp58/Rp58 tetramer, in the PDB and BFMD databases (24). Despite the lack of templates and very low number of orthologous sequences available to AlphaFold for structure prediction (<30), the predictions generally have high confidence scores, especially in structured interface regions (average ɑ-helix pLDDT ≈90) (**Table S1**, **supplementary information**).

### Prediction of Mp1/Mp58 covarying residues clusters

Exploratory covariation analysis of species-matched Mp1 and Mp58 alignments identified multiple high-ranking cross-protein residue pairs. Candidate pairs were ranked by average-product-corrected mutual information (MI-APC), and empirical support for the highest-ranking pairs was assessed by one-sided permutation testing of the corresponding raw MI values using 5,000 permutations per tested pair. The strongest residue-identity covariation signals involved Mp1 F25, V32 and the I43–R47 region, together with Mp58 G22, D30 and T34, with residue positions reported relative to the mature proteins following signal-peptide removal. Additional high-ranking signals involved Mp1 K72 and K83 and Mp58 K61, Q72, K75 and A76.

In the residue-identity analysis, each amino acid was treated as a distinct categorical state irrespective of physicochemical similarity; consequently, no similarity-based weighting was applied to conservative versus non-conservative substitutions. A complementary physicochemical-class analysis, in which residues were grouped according to aromatic, hydrophobic, positively charged, negatively charged, polar uncharged, or special/structural properties, identified additional candidate associations based on conservation and covariation of broader residue properties (**Figure S6**). As these signals were not dominant in the residue-identity analysis, they were treated as exploratory candidates for coordinated variation in residue properties. Integration of both sets of analyses identified three broader clusters (**Figure 4A, 4B, supplementary information**). The primary interface-associated cluster comprises Mp1 F25, V32 and I43–R47 with Mp58 G22, D30 and T34; a secondary basic/property-associated cluster comprisese Mp1 K72, K75 and K83 with Mp58 K61, Q72, K75, A76, D90, K105 and D106; and a C-terminal/property-associated extension comprises Mp1 E27, S54, L62, I66, D71, M85 and H89 with Mp58 L116, V123, A124 and Q125, overlapping the secondary cluster at Mp58 K105, D106 and V123 (**Figure 4A, 4B, S6)**

**Figure 4.**
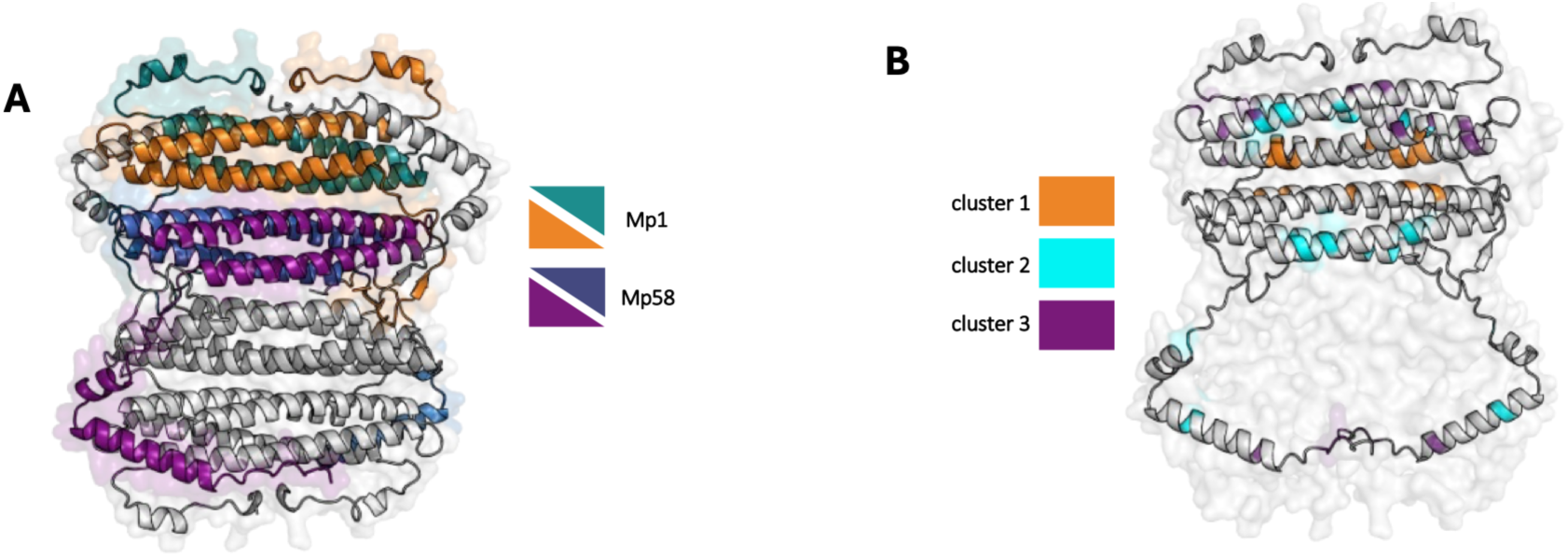
Clusters of co-variant residues based on Mp58 and Mp1 protein sequence alignments. **(A)** AlphaFold3 Mp1+Mp58 hetero-octamer (4+4), with one half coloured by subunit. Mp1 is shown in teal/orange and Mp58 in blue/purple; the symmetry-related half is shown in grey. **(B)** The same oligomer shown as a faint surface for the lower half, with the upper Mp1+Mp58 chains in grey and covariation clusters highlighted. Cluster 1 (orange) comprises the primary Mp1-Mp58 interface centred on Mp1 R47 and Mp58 D30/T34; cluster 2 (teal), comprises a secondary electrostatic/property-associated region; and cluster 3 (purple) comprises a C-terminal/property-associated extension.

Mapping the covariation-associated residues onto the three highest-ranked AlphaFold models showed that recurrent direct Mp1-Mp58 contacts were concentrated within cluster 1 (**Figure 4B**). Mp1 R47 formed contacts with Mp58 D30 and T34, which were retained across models. The R47-D30 interaction showed either salt-bridge, van der Waals and hydrogen bonding across models and is therefore best described as a recurrent charge-compatible contact rather than a uniformly resolved salt bridge. Other cluster 1 residues did not consistently form explicit cross-chain contacts but were positioned within or adjacent to the same predicted interface. These residues may contribute indirectly by maintaining local packing, side-chain chemistry or backbone geometry around the R47-D30/T34 contact region. Their covariation is therefore consistent with preservation of local interface architecture, although this remains an inference rather than direct evidence of residue-specific co-evolution. Clusters 2 and 3 did not form recurrent direct Mp1-Mp58 interfaces in the analysed models. Cluster 2 was dominated by intrachain contacts consistent with roles in maintaining local electrostatic or structural stability. Cluster 3 was likewise dominated by local contacts within Mp1 or Mp58.

We attempted to disrupt the Mp1-Mp58 interaction based on RING or PDBePISA, which identify the minimal number of interface side chains contributing important stabilizing interactions in the AlphaFold models. However, the Mp1-Mp58 interface differed from contacts within the Mp58 tetramer, with less prevalent single hydrophobic interactions. Alanine substitution of three selected residues simultaneously in Mp1-Mp58 (Mp58 Y19, Mp58 D30, Mp1 F40) did not detectably reduce interaction in *E. coli* lysates (**Figure S7**).

### Mp1/Rp1 evolved different oligomerization states

To determine the oligomeric state of Mp1/Rp1 in the absence of Mp58/Rp58, we produced recombinant Mp1 or Rp1 fused to maltose binding protein (MBP) in *E. coli* and investigated mass by SEC-MALS. Surprisingly, we measured a mass of ≈50 kDa (monomer) for MBP-Mp1 and a mass of ≈220 kDa (tetramer) for MBP-Rp1, pointing to different oligomerization states for Mp1 and Rp1 (**Figure 5A**). AlphaFold structure predictions of Mp1 and Rp1 tetramers have high confidence scores, especially the prediction of the Rp1 core when protruding loops are removed (pTM = 0.92) (**Table S1**). The core of the predicted tetramer has a different pattern of residue interactions in Rp1 compared to Mp1. While 2 disulfide bonds appear at the center of Rp1 (Cys43), Mp1 contains 0 cysteine residues (**Figure 5B, 5C**). At this position, ≈60 % of the aligned orthologous sequences contain a cysteine and ≈40 % contain isoleucine, with no other amino acids occurring except valine once (**Figure S6**). However, incubation of Rp1 in strong reducing conditions or mutation of this cysteine to alanine did not cause a change in the Rp1 oligomerization state, indicating that other residues contribute to tetramer stabilisation (**Figure S8**). There is a notable structural relationship between homo-oligomeric Rp1 and the hetero-oligomeric complex in that the structure comprised of two dimers of each protein (2x Mp58 + 2x Mp1, or 2x Rp58 + 2x Rp1) has the same conformation as the predicted structure of Rp1 tetramer, as reflected by high TM scores (>0.90) (**Figure 5D**, **S2**).

**Figure 5.**
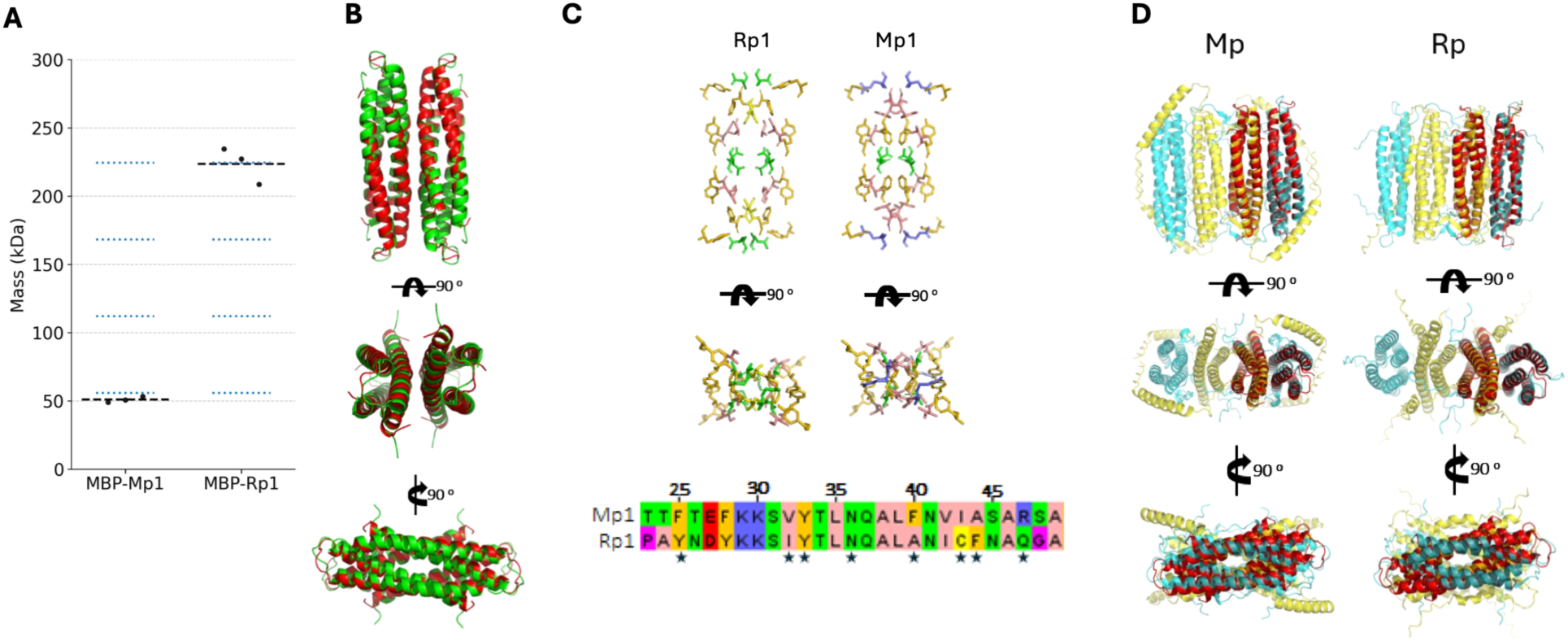
Structure of aphid effectors Mp1/Rp1. **(A)** Size exclusion chromatography with multi-angle light scattering (SEC-MALS) results for maltose binding protein (MBP)-tagged Mp1 and Rp1 proteins, expressed in and purified from *E. coli*. Graphs shows mass measured for 3 biological replicates (black dots), mean mass (black dashed line), and expected mass of oligomerization states 1-5 (blue dashed lines). **(B)** AlphaFold3 tetrameric models superimposed using PyMOL “super” command. Rp1 is shown in red, Mp1 is shown in green. Models are shown in PyMOL cartoon style in three orientations rotated by 90°. **(C)** amino acid residues at the core of Rp1 and Mp1 tetramers displayed in **B**, shown in PyMOL stick style, coloured by amino acid side chain chemical property (Zappo scheme). The sequence alignment with corresponding colours is shown below, displayed residues are marked with a star symbol. **(D)** same as **B**, using different AlphaFold3 models. Tetrameric Rp1 is shown in red superimposed on the octameric AlphaFold3 prediction of either the *Myzus persicae* (Mp) or *Rhopalosiphum padi* (Rp) effector complex with 4 Mp58/Rp58 (yellow) and 4 Mp1/Rp1 (cyan) subunits.

### Mp58 oligomerization is required for Mp1-Mp58 complex formation and effector activity

Inspection of the Mp58 and Rp58 tetramer structures suggested that a conserved leucine or phenylalanine residue at position 67 (Mp58^F67^, Rp58^L67^) holds together two dimers by hydrophobic interactions (**Figure 6A**). To test this, we mutated this residue to glutamate or alanine in both Mp58 and Rp58, which led to a pronounced shift in the SEC elution peak to a higher elution volume (**Figure 6B, S9A**). We compared this data to the SEC calibration graph, using the peak apex elution volume of standard proteins of known mass to construct a line of best fit, and plotting the data for full-length and chymotrypsin-digested Mp58/Rp58 tetramers, all of which had mass measurement from SEC-MALS available (**Figure S9A**). This showed that the observed elution volume of Mp58/Rp58^F/L67^ mutants likely correspond to a dimer with a similar volume/mass ratio, rather than a tetramer in a more compact conformation (**Figure S9B**). In the case of Mp58^F67A^, a mixture of tetramer and dimer peaks appeared after the protein was stored at 4 °C, while Rp58^L67A^ remained dimeric only (**Figure S10A**). Further analyses identified a conserved tryptophan residue at the monomer-monomer interface within each dimer (W66), which we predicted was important for dimerization. Mutation of this residue to glutamate or alanine resulted in a slightly higher main peak elution volume, which may be indicative of a more compact dimer conformation (**Figures 6B, S9B, S10B**). Nano differential scanning fluorimetry (nanoDSF) of Mp58 and Rp58 showed a high unfolding temperature (>70 ℃) of the intra-dimer interface, with W66 being the main source of signal as the only tryptophan residue in the proteins (**Figure S11**). This suggests that monomers are thermodynamically unfavorable. Overall, these results show that Mp58 and Rp58 have evolved the same primary oligomerization state (tetramer), with different potential to convert into lower oligomerization states and perhaps different conformations under *in vitro* conditions.

**Figure 6.**
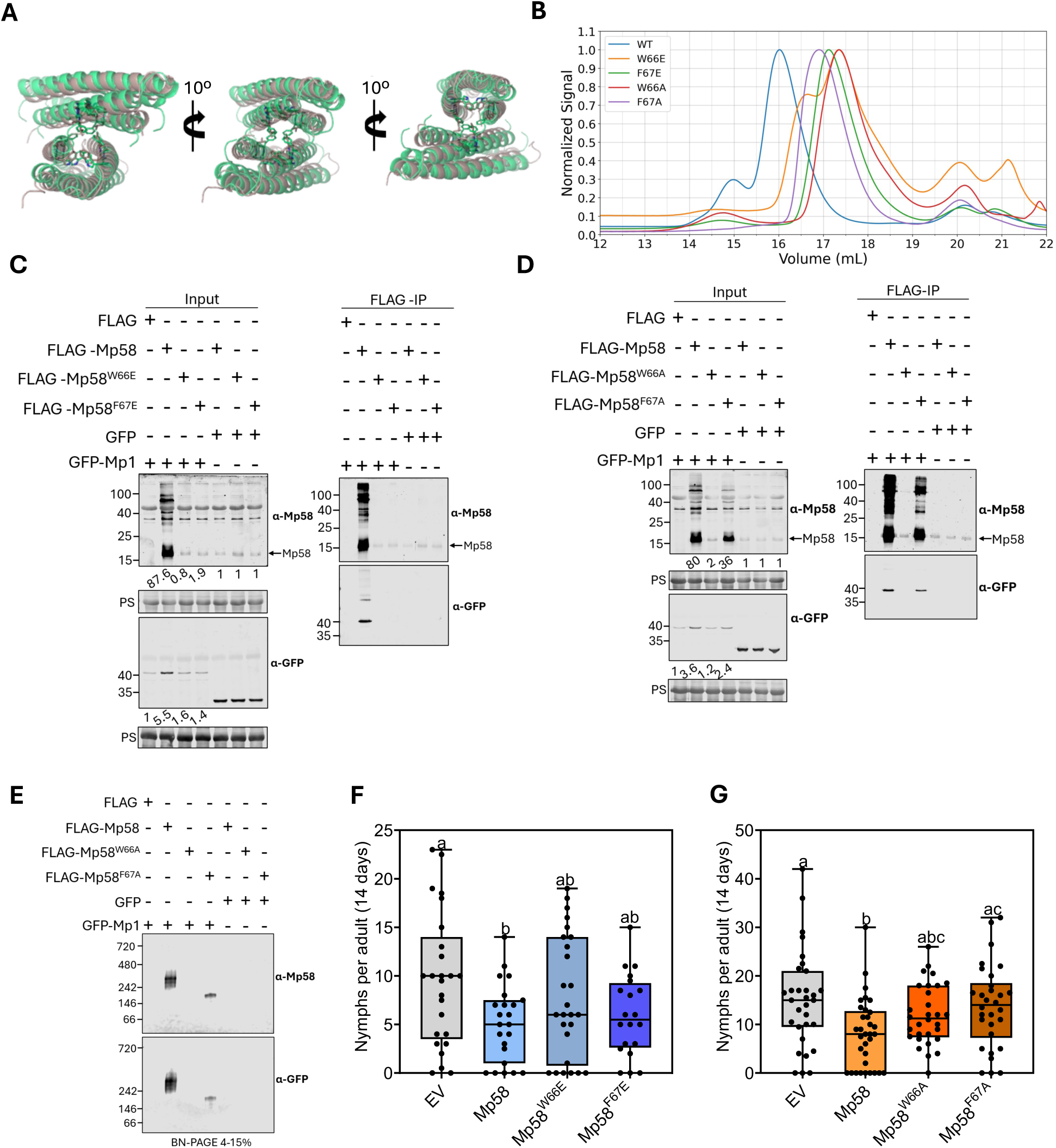
Mp58 oligomerization is required for Mp1 interaction and effector activity. **(A)** Protein models superimposed using PyMOL “super” command. The models show the Rp58 chymotrypsin fragment crystal structure (brown) and tetrameric Mp58_3-83_ AlphaFold 3 prediction (green). Models are shown in PyMOL cartoon style in three orientations rotated by 10°; residues with sequence indices 66 and 67 are shown in stick representation. **(B)** Size exclusion ultraviolet chromatograms of Mp58 sequence variants. **(C)** Co-immunoprecipitation assay of Mp58 glutamate substitution mutants with Mp1. Sodium dodecyl sulfate polyacrylamide gel electrophoresis (SDS-PAGE) and immunoblotting (IB) was used before (input) or after immunoprecipitation using FLAG beads (FLAG-IP). Numbers refer to mass of PAGE marker proteins (kilodaltons, left of image) or band intensity of FLAG-Mp1/Mp58 and GFP-Mp1/Mp58 normalized to total protein (under image). Quantitation as performed using Empiria Studio and LI-COR images. FLAG-Mp58 variant levels in samples co-expressed with GFP-Mp1 were compared to protein levels in samples co-expressed with GFP (control, set at 1) to generate band intensity ratios; GFP-Mp1 levels in samples co-expressed with Flag-Mp58 variants were compared to protein levels in samples co-expressed with Flag (control, set at 1) to generate band intensity ratios. PS indicates Ponceau Stained blots showing loading and transfer. **(D)** same as **C**, but using Mp58 alanine substitution mutants. **(E)** blue native (BN-PAGE) of FLAG-IP samples used for SDS-PAGE in **D**. For C, D and E data is representative of 2-3 independent replicates. **(F G G)** *Myzus persicae* fecundity assay on *Nicotiana benthamiana* leaves expressing Mp58 glutamate **(F)** and alanine **(G)** variants; means that do not share a letter are significantly different (*P* < 0.05, one-way ANOVA with Tukey *post-hoc* test). Data are representative of n=3 independent biological replicates with n=6-10 per independent replicate for data in F and n=9-12 per independent replicate for data in G.

To assess effector complex formation and oligomerization *in planta*, we generated plant expression constructs for Mp58 where the conserved tryptophan or phenylalanine residues at positions 66 and 67 were mutated to either glutamate (Mp58^W66E^/Mp58^F67E^) or alanine (Mp58^W66A^ / Mp58^F67A^). All Mp58 mutants were detected upon transient expression in *N. benthamiana* by immunoblotting at comparable levels (**Figure 6C, 6D**). We determined whether the Mp58 mutants were able to interact with Mp1 by co-immunoprecipitation of co-expressed FLAG- and GFP-tagged effectors. We observed an increase in detectable FLAG-Mp58 expression in the presence of GFP-Mp1 compared to the GFP control, but no similar increase was detectable for FLAG-Mp58^W66E^or FLAG-Mp58^F67E^ (**Figure 6C, 6D**). In line with this, GFP-Mp1 co-immunoprecipitated with FLAG-Mp58 but not with FLAG-Mp58^W66E^ or FLAG-Mp58^F67E^, indicating that these mutants lost the ability to interact with Mp1 (**Figure 6B**). In similar experiments with Mp58 mutants, where we substituted with alanine, we only observed an increase in FLAG-Mp58^F67A^, but not FLAG-Mp58^W66A^, levels in the presence of GFP-Mp1, although this increase was less pronounced when compared to FLAG-Mp58. In line with this, GFP-Mp1 co-immunoprecipitated with FLAG-Mp58^F67A^, but not with FLAG-Mp58^W66A^ (**Figure 6D**). BN-PAGE followed by immunoblotting of the same samples showed a faster migrating band corresponding to the Mp58^F67A^-Mp1 complex compared to a band corresponding to Mp58-Mp1, indicating a change in the oligomerization state of the mutant complex. No band was detected for FLAG-Mp58^W66A^, in line with the observation that this mutant does not interact with or is stabilized by GFP-Mp1 (**Figure 6D**). These results indicate that in a potential monomeric state (Mp58^W66E/A^), Mp58 can no longer form a complex with Mp1 *in planta*. When Mp58 is present as a dimer, Mp58 is still able to form a complex with Mp1 (Mp58^F67A^), however this can be disrupted by creating a negative charge (Mp58^F67E^) around the interacting region of the complex.

To determine whether Mp58 oligomerization is required for effector activity, we performed *M. persicae* performance assays on *N. benthamiana* leaves transiently expressing Mp58 or Mp58 mutants. In line with previous work (18), aphid fecundity was reduced on infiltration sites expressing Mp58 with a ±40-50% reduction in the mean number of nymphs produced per adult aphid compared to the vector control in the glutamate and alanine mutant assays (**Figure 6E, 6F**). In contrast, aphid fecundity was not significantly different on leaf areas expressing Mp58 W66 and F67 mutants when compared to those expressing the vector control (**Figure 6E, 6F**). Therefore, the disruption of Mp58 oligomerization correlates with a loss of its effector activity in decreasing host susceptibility to aphids.

## Discussion

Plant pathogenic microbes and herbivorous insect pests produce effectors with a wide range of activities in modulating host cell responses. Interestingly, these effectors may form hetero-oligomeric complexes that are predicted to regulate effector translocation and activity. The mechanisms underlying such complex formation and its biological significance are largely unknown. Here, we show that an effector pair, encoded by genetically linked genes conserved across aphid species, form a hetero-hexameric/octameric complex that is structurally distinct from any currently available structures in the PDB. We show that hetero-oligomeric effector complex formation leads to stabilisation of the effectors and identify critical residues at the interaction interface. Disruption of the interaction interface through mutation of single residues not only prevents hetero-oligomeric complex formation but also reduces effector activity *in planta*. Our data point to a model wherein reversible transitions between effector-effector self-assemblies and effector-host target complexes may influence effector functionality at the host interface.

Despite low sequence alignment depth for both Mp1/Rp1 and Mp58/Rp58, and the lack of structural/sequence templates in databases, the Alphafold3 prediction of the core Mp58/Rp58 tetramer within the effector complex was supported by high confidence scores and validated by the crystal structure. This accuracy may be due to a high level of conservation amongst a small number of orthologues, and/or the strong physical stability of the structure. For example, it is known that local physical environments (*e.g.* hydrophobic core packing) can drive high prediction confidence and accuracy independently of co-evolution when sequence alignment depth is greatly reduced (25, 26). Notably, the Alphafold3 prediction for the Mp58+Mp1 or Rp58+Rp1 tetrameric subcomplex as well as the Mp1/Rp1 tetramer were supported by high confidence scores and significant amino acid conservation. However, further experimental validation through amino acid substitutions is needed to confirm Mp1/Rp1 substructures. Strikingly, a single mutation at the inter-dimer or intra-dimer interface in Mp58/Rp58 can readily convert this tetramer into a dimer with significant thermal stability. These results suggest a model in which the effector complex is viewed as an array of juxtaposed homodimers, with the potential to dissociate into isolated dimers and allow for subunit exchanges (*e.g.* Rp1 monomers exchanging between the Rp1 homotetramer and the Rp58-Rp1 heterotetramer). The interdependence in oligomeric assembly is emphasized by the observation that disruptive mutations in the tetramer, notably those involving glutamate substitution or intra-dimer interface disruption, resulted in lower mobility in SEC and prevented Mp1 interaction *in planta*, while the Mp58 F67A mutant retained the ability to interact with Mp1 and form a tetramer. We propose that this effector complex may be dynamic, possibly relating to its function in regulating other effector proteins and or/interacting with target plant proteins.

The sequence alignments for the two proteins in the effector complex comprise a relatively small number of sequences (20–30), and no similar sequences are found outside the *Aphididae* family. This suggests that the coevolutionary arms race between plants and aphids led to a relatively isolated branch within the evolutionary tree of proteins. The conserved, head-to-tail tandem architecture and high promoter homology observed between these two aphid-specific salivary effectors support a birth-and-death model of gene duplication followed by neofunctionalization (27, 28). The similarity in promoter regions (27) suggests strict evolutionary constraints to preserve stoichiometric balance, in line with our observed 1:1 ratio for effector complex assembly. It is possible that one or both proteins evolved to form tetrameric assemblies either before or shortly after a gene duplication event. Following this duplication, sequence divergence between the genes may have led to subtle conformational differences in the assembly of monomers or dimers and the evolution of a novel heteromeric complex, transforming a homomeric or monomeric ancestral protein into a two-component aphid-specific effector complex with a functional advantage. How heteromeric complex formation is associated with potential diversification of host target interactions remains an open and important question. While Mp1 and Mp58 associate with host cell trafficking proteins VPS52 in host plants, we have not found evidence that orthologs Rp1 and Rp58 share this target in the *R. padi* host barley (16, 17). Further characterization of these effectors and the complexes they form across aphid-host systems will be needed to gain further insights into how this pair co-evolved with different host plant species. Nonetheless, the conservation of the effector pair across the aphid evolutionary tree and the formation of a hetero-oligomeric complex highlight that these effectors function as a unit that is significant during plant-aphid interactions.

Co-variant analyses alongside structural modelling and residue contact analysis showed that co-varying residues between Mp1 and Mp58 were identified outside of the primary structural interface between the two proteins. We hypothesise that these residues may contribute to the stabilisation of the proteins upon complex formation or may be involved in interactions with shared target proteins, such as AtVPS52. Future work, based on molecular dynamics (MD) simulations, could help further resolve the structural dynamics of the Mp1-Mp58 effector complex, ideally in the context of host target interactions. In this study, we omit MD because the potential subunit exchanges and dissociations that we hypothesise may occur in this complex would require a large solvation shell, which, coupled with the large complex size (>120 kDa), would require significant computational resources. Alternatively, structural dynamics could be explored through cryo-electron microscopy, which has the potential to capture different conformations that may shed light on how the structure relates to the functional properties of the complex.

While effector oligomerization occurs in host-pathogen interactions more widely and may be involved in effector translocation and/or regulation, its functional importance to plant pathogenic microbes and pests is yet to be fully explored. Our work shows that detailed effector characterization, including at the structural level, may unveil novel protein folds as well as oligomeric assemblies at the plant-pathogen/pest interface. In addition to the Mp1-Mp58 effector pair, aphid genomes feature additional conserved, co-located and co-expressed candidate effector pairs, and the Mp1-Mp58 pair is tightly co-regulated with a subset of aphid salivary proteins (27). This raises the possibility that, in addition to Mp1-Mp58, other effectors may be involved in complex formation as part of an effective virulence strategy. On the plant side, it is well established that the receptors that detect the presence of effectors, nucleotide-binding leucine-rich repeat receptors (NLRs), form larger oligomeric complexes, called resistosomes, to mediate an immune response (29). For example, ZAR1, in the presence of bacterial effectors, induces the formation of a hetero-oligomeric protein complex, which acts as a calcium channel in the plant immune response (30–32). This NLR oligomerization strategy is widespread in plants and considered a central mechanism underlying plant resistance to pathogens (29). However, protein oligomerization at the plant-pathogen/pest interface does not appear to be unique to plant proteins and can, at least in some cases, extend to pathogen or pest effector proteins. Overall, our findings highlight the importance of effector hetero-oligomerization at the interaction between plants and herbivorous insects, supporting the model that this effector pair represents an evolutionarily conserved module among different aphid species.

## Materials and Methods

### Phylogenetic Tree Generation

Phylogenetic analyses were performed independently for Mp1 and Mp58 homologue sequences identified in aphid species. Protein sequences were screened for N-terminal signal peptides using SignalP 6.0 (33) and predicted signal peptide regions removed prior to orthology assignment using OrthoDB v12.2 (34). and subsequent alignment. For Mp58, no signal peptide was predicted for Tt58 and therefore no signal peptide region was removed from this sequence.

The 28 homologous amino-acid sequences from each effector family were aligned with Phylogeny.fr (35) using MAFFT v7.467 (36) with the L-INS-i algorithm, incorporating both local pairwise alignment information and iterative refinement. Alignments were created using the BLOSUM62 scoring matrix (37), a gap opening penalty of 1.53 and offset value of 0.123. Ambiguously aligned regions were removed using BMGE (38) with the BLOSUM30 similarity matrix, maximum entropy threshold of 0.5, gap-rate cut-off of 0.5 and sliding window size of 3. The Mp58 alignment contained 28 taxa and 116 amino-acid positions; the Mp1 alignment contained 28 taxa and 106 amino-acid positions. Maximum-likelihood phylogenetic trees were inferred using PhyML v3.3.20190909 (39) under the LG amino-acid substitution model. Amino-acid equilibrium frequencies were model-derived, starting topologies generated using BioNJ (40), branch lengths and substitution model parameters optimised and among-site rate heterogeneity modelled using a discrete gamma distribution with four rate categories. Invariant sites and gamma shape parameter were estimated (0.015 and 1.548 respectively for Mp58; 0.000 and 1.749 respectively for Mp1). Tree topology searching was performed using subtree pruning and regrafting, with the best topology retained and branch support assessed using 1,000 bootstrap replicates for each tree. Bootstrap support was summarised using BOOSTER (41) and normalized transfer bootstrap expectation support values mapped onto final maximum-likelihood topologies.

### Comparison of Phylogenies

The maximum-likelihood trees inferred for each effector family were compared using a topology-based approach with Visual TreeCmp (42) in overlapping pair comparison mode. Trees were pruned to shared taxa and compared using Robinson-Foulds (RF) distance as a measure of shared internal split structure. Quartet distance was also calculated as an additional measure of topological congruence based on conservation of four-taxon relationships. Normalised distances and summary statistics were exported from Visual TreeCmp.

Testing of potential correlation of evolutionary distances between Mp1 and Mp58 was conducted using pairwise patristic distance matrices calculated independently from the Mp1 and Mp58 maximum-likelihood trees. This analysis was conducted to determine if taxa separated by greater evolutionary distance in the Mp1 phylogeny also tended to be separated by greater evolutionary distance in the Mp58 phylogeny. Correlation between patristic distance matrices was assessed using a Mantel permutation test with Spearman correlation and 9,999 permutations since pairwise patristic distances are non-independent.

### Inter-Protein Residue Covariation Analysis

Inter-protein residue covariation between Mp1 and Mp58 was analysed using custom Python scripts executed in Spyder 5.5.1 under Python 3.12.4 (Anaconda distribution), using NumPy 1.26.4 and pandas 2.2.2. Independently aligned, BMGE-trimmed Mp1 and Mp58 protein alignments were matched by taxon using shared sequence identifiers, and only taxa represented in both protein families were retained. Residue numbering was referenced to the *M. persicae* sequences. Covariation was evaluated exclusively between Mp1 and Mp58 alignment columns; within-protein Mp1-Mp1 and Mp58-Mp58 column pairs were not considered.

Two parallel analyses were performed. In the residue-identity analysis, each of the 20 standard amino acids was treated as a distinct categorical state. In the physicochemical analysis, amino acids were grouped into six classes: aromatic (F, Y, W, H), aliphatic hydrophobic (A, V, I, L, M), positively charged (K, R), negatively charged (D, E), polar uncharged (S, T, N, Q), and special/structural (G, P, C). Alignment gaps, ambiguous residues and non-standard amino-acid symbols were treated as missing states.

For each analysis, alignment columns were retained if at least 50% of paired sequences contained a non-gap state and at least two distinct non-gap encoded states were present. The variability criterion was therefore applied to individual amino-acid identities in the residue-identity analysis and to physicochemical classes in the physicochemical analysis. Mutual information (MI) was calculated for all retained Mp1–Mp58 column pairs using base-2 logarithms, with taxa containing a missing state at either member of an individual pair excluded from that pairwise calculation.

The resulting cross-protein MI matrix was subjected to average product correction (APC), and candidate covarying pairs were ranked according to APC-corrected MI (MI-APC). APC was used to reduce broad background MI associated with columns showing generally elevated covariation across multiple positions and was not treated as an explicit correction for phylogenetic non-independence. Empirical support for the 100 highest-ranking MI-APC pairs, or all pairs where fewer than 100 were available, was assessed using one-sided permutation tests of the corresponding raw MI values. For each pair, the Mp58 state vector was randomly permuted relative to Mp1, disrupting the original taxon pairing while retaining the overall composition of the permuted column before pairwise exclusion of missing observations. A total of 5,000 permutations was performed per pair using a fixed base random seed of 12345, with a deterministic rank-specific seed used for each candidate pair. Empirical P values were calculated using an add-one correction, (k+1)/(n+1), where (k) is the number of valid permutations producing an MI greater than or equal to the observed MI and (n) is the number of valid permutations.

Because the number of species-matched sequences was limited and phylogenetic non-independence was not explicitly modelled, the analyses were treated as exploratory. Permutation P values were not corrected for multiple testing, and covariation signals were therefore interpreted as candidate associations in conjunction with residue conservation and structural interaction analyses rather than as independent evidence of direct residue coupling. Analysis scripts are available at [https://github.com/SRF-Boltz/Covariation_Analysis/releases/tag/V1.0.0].

### Protein structure prediction and analysis

Protein structures were predicted using the AlphaFold 3 web server (43). Five structural predictions were generated in each case and ranked using AlphaFold confidence metrics. Model quality was assessed using the overall ranking score, predicted template modelling score (pTM), interface predicted template modelling score (ipTM), predicted local distance difference test (pLDDT), predicted aligned error (PAE), and the presence or absence of steric clashes. The top-ranked model was used for most analysis, while protein interface detail was assessed cautiously by comparing residue contact patterns across the 3 highest-ranked models. Confidence in local structural regions was visualised by colouring the model according to pLDDT with PyMol (“The PyMOL Molecular Graphics System, Version 3.1 Schrödinger, LLC”), and inter-chain/domain placement confidence was evaluated using the PAE plot.

The average pLDDT score of ɑ-helical regions of the models was calculated from the B factor column of PDB files. Valdar conservation score was calculated from sequence alignment using Jalview (44), and inserted into the B factor column for visualization in PyMOL. The Zappo amino acid color scheme was applied to protein structure models by customised PyMOL script. Code is available (**supplementary information).** To identify residue candidates for mutation, protein structure models were submitted to RING (45) or PDBePISA (46) using default settings.

To assess differences in oligomer conformation between protein structure models, the top ranked AlphaFold 3 models were processed. Only tetrameric models of similar size were used - octamer models were converted to tetramer using PyMOL to select the four chains comprising the centre or one half of the octamer. All vs all structural alignment of models was performed using custom python code to run MMalign in the USalign package (47, 48) (command line options -mm 1 -ter 0) or PyMOL “super” command with custom-coded standard TM score calculation method. Custom python code (**supplementary information**) was used to visually inspect the alignments and generate TM score heatmaps from the determined matrices.

### Plasmid cloning

To generate plant expression constructs, Mp1 and Mp58 were cloned into pDONR207 (Gateway entry vector) using BP Clonase II (Thermo Fisher Scientific, 11789020) following the manufacturer’s instructions. To generate Mp58 variants, non-overlapping primers were designed using NEBase Changer (**supplementary information**) to amplify the full-length plasmid, using Mp58 pDONR207 as the template. Following amplification, linear PCR products were circularised by incubating with 50 ng PCR product with 1X T4 DNA Ligase Reaction Buffer (Thermo Fisher Scientific, B69), 1 µl T4 DNA ligase (Thermo Fisher Scientific, EL0011), and 1 µl Dpn1 (New England Biolabs, R0176S), and 1 µl T4 Polynucleotide Kinase (New England Biolabs, M0201S) in a 10 µl reaction for 1 hour at room temperature (RT). Inserts were transferred into Gateway destination vectors using LR clonase II (Thermo Fisher Scientific, 11791020) following the manufacturer’s instructions. pEarleyGate 202 (FLAG tag) and pB7WG2 (no tag) destination vectors were used. Green fluorescent protein (GFP) tagged Mp1/Rp1 and Mp58/Rp58 were cloned and described previously (17, 18). All plasmids were transformed into *Agrobacterium tumefaciens* for expression in *Nicotiana benthamiana*.

For *E. coli* protein expression and purification, codon-optimized maltose binding protein (MBP), Mp1, Rp1, Mp58 and Rp58 coding DNA sequences were synthesized. Regions of interest were PCR-amplified and assembled into a pET-15b expression vector (**supplementary information**) using GeneArt Gibson Assembly HiFi Master Mix (Thermo Fisher Scientific). 6×His tags. tobacco etch virus (TEV) or Human Rhinovirus 3C (3C) protease cleavage sites and mutations were encoded in PCR primers (**supplementary information**).

All circularised constructs were transformed into Top10 cells (Thermo Fisher Scientific, C404003) and verified by sequencing.

### Polyacrylamide gel electrophoresis (PAGE)

For sodium dodecyl sulfate PAGE (SDS-PAGE), protein samples were mixed into protein loading buffer (Li-Cor, 928-40004) or Laemmli buffer [50 mM Tris-HCl (pH 6.8), 10 % glycerol, 2 % SDS, 0.05 % Orange G, 100 mM DTT]. Samples were incubated at 80-90 °C for 5-10 min, loaded on 4-20 % PROTEAN® TGX™ Precast Protein Gels (Bio-Rad, 4561084) or homemade 15 % resolving SDS-PAGE gel, and run at 100–200 V until the dye reached the bottom of the gel. PageRuler™ Prestained Protein Ladder (Thermo Fisher Scientific, 26616) was loaded to identify approximate sizes of proteins. Prior to immunoblotting, proteins were transferred immediately after PAGE to either nitrocellulose or polyvinylidene difluoride (PVDF) membranes, using a Tris–glycine transfer buffer (25 mM Tris, 192 mM glycine, 20% ethanol) for 90–120 min at 90 V or using the Bio-Rad Turbo blot system following manufacturer’s instructions.

Blue native PAGE (BN-PAGE) was adapted from (49). Protein extracts and immunoprecipitation (IP) eluates were added with 4× NativePAGE™ Sample Buffer (Invitrogen™, BN2003) and NativePAGE™ 5 % G-250 Sample Additive (Invitrogen™, BN2004) or a homemade version (5 % w/v Coomassie G-250 in 500 mM 6-aminocaproic acid) to a final concentration of 0.125 %. BN-PAGE was run using the Bio-Rad Mini-Protean gel system. Samples were loaded on Bio-Rad 4-15 % or 4-20 % PROTEAN® TGX™ Precast Protein Gels (Bio-Rad, 4561084) and run in cathode buffer containing Coomassie G-250 (by adding NativePAGE™ Cathode Buffer Additive to 1/200 dilution, Invitrogen™ to Tris/Glycine running buffer) 100 V for 1/3rd of the gel and 150 V until the dye front reached the bottom of the gel. NativeMark™ Unstained Protein Standard (Invitrogen™, LC0725) was loaded to identify approximate sizes of proteins. Proteins were transferred to PVDF membranes in Towbin buffer with 10 % EtOH and 0.01 % SDS.

### Total protein staining

For in-gel total protein detection, SDS-PAGE gels were stained using Imperial™ Protein Stain (Thermo Fisher Scientific, 24615) or Bio-Safe™ Coomassie Stain (Bio-Rad, 1610786) following manufacturer’s instructions. For total protein detection on membranes, the protocol was based on LI-COR instructions. Membranes were dried at 37 ℃ or RT, rehydrated with ethanol (PVDF membranes) or PBS (nitrocellulose membranes) and stained with Revert™ 700 Total Protein Stain (Li-Cor, 926-11021) or a homemade version [30% ethanol, 7% acetic acid, 0.001% Fast Green FCF (Thermo Fisher Scientific, A16250)] for 5-10 min. Membranes were washed with a wash solution (30% ethanol, 7% acetic acid) to remove background and imaged using a Li-Cor odyssey CLx with the 700 nm channel. The stain was removed with a destain solution (30% ethanol, 100 mM sodium hydroxide). Membranes were rinsed with PBS or dH_2_O before proceeding to blocking.

### Immunoblotting

When analyzing protein samples derived from *E. coli*, membranes were blocked with 2 % skimmed milk powder in PBS-T (phosphate-buffered saline containing 0.1 % Tween-20) before incubating with primary antibodies overnight at 4 °C or 1 hour at RT. The following antibodies were used: his (Thermo Fisher Scientific, MA1-21315), strep (Merck, MAC143). When analyzing protein samples not derived from *E. coli*, membranes were blocked with 5 % skimmed milk powder in TBS-T (Tris-buffered saline containing 0.2 % Tween-20) before incubating with primary antibodies overnight at 4 °C. The following antibodies were used:

FLAG (Protein Tech, 20543-1-AP), GFP (Santa Cruz Biotechnology, sc-9996), native Mp58 antibody. Membranes were washed with PBS-T or TBS-T before incubating with respective secondary antibodies, mouse (LI-COR, 926-33210) or rabbit (LI-COR, 926-32211) in the dark at RT for 1 hour. Proteins were detected with LI-COR Odyssey CLx in the 700 and 800 nm channels.

Protein quantification was conducted by normalizing the band intensity of indicated proteins against the total protein amounts using Empiria Studio v.3.3. Relative ratios of signal intensity within experimental set-ups were calculated based on comparisons to relevant control samples.

### *Myzus persicae* saliva collection

*M. persicae* (clone O) were maintained in a greenhouse on *Solanum tuberosum* (cv Desiree). Approximately 100 mixed aged aphids were placed in plastic pots with the bottom removed and sealed with gauze. The top was sealed with two stretched sheets of parafilm holding 300 µl filter sterilised 10 % sucrose. The aphids were reared on the diets at 20 °C with 18 hours light and 6 hours dark and 65 % relative humidity for 24 hours. Diets were pooled (1.2 ml from ≈400 aphids), snap-frozen in liquid nitrogen, and stored at -80 °C until use. Pooled saliva samples were concentrated to ≈50 µl using Pierce™ Protein Concentrators with 3 kDa MWCO (Thermo Fisher, 88515) at 4 °C. The concentrated sample was then split in half between SDS-PAGE and BN-PAGE.

### *Myzus persicae* performance assays on *Nicotiana benthamiana*

Three-week-old *N. benthamiana* plants were infiltrated with the indicated with *A. tumefaciens* carrying the indicated constructs. Two adult aphids were placed on the underside of leaves 1 d after infiltration and contained with clip cages. The following day, adults were removed, and two 1st instar nymphs were left on each infiltration site. At 7 d post-infiltration, nymphs were transferred to new agroinfiltrated leaves and aphids were counted 7 d later (14 d post-initial agroinfiltration).

### *Agrobacterium tumefaciens* infiltration for transient expression

*A. tumefaciens* (strain GV3101) carrying the indicated constructs were grown shaking overnight at 28 °C. Cultures were centrifuged at 2500 g for 10 min at RT. Pellets were resuspended in infiltration buffer (10 mM MgCl_2_ and 10 mM MES, pH 5.6), the OD_600_ was adjusted and 200 µM acetosyringone was added to the cell suspension. For co-IP experiments, cell suspensions were adjusted to OD_600_=0.3 for all constructs except GFP (OD_600_=0.02) and p19 (OD_600_=0.1); for aphid performance assays, OD_600_=0.2 was used for all constructs except p19 (OD_600_=0.1). Cultures were incubated for 2 hours in the dark at 28 °C. Three-to four-week-old *N. benthamiana* plants were infiltrated with cell suspensions carrying the indicated constructs.

### Co-immunoprecipitation

Leaf tissue from *N. benthamiana* (12 leaf discs, 1.5 cm) was ground in liquid nitrogen and protein was extracted with cold GTEN buffer (10 % glycerol, 25 mM Tris–HCl pH 7.5, 1 mM EDTA, 150 mM NaCl) containing 0.2 % Triton X-100, 1 mM dithiothreitol (DTT) and 1× EDTA-free protease inhibitor cocktail (Thermo Fisher Scientific, A32965). Samples were incubated on ice for 10 min with frequent vortexing. Samples were centrifuged at 16000 g for 20 min at 4 °C. The supernatant was diluted with cold IP buffer (GTEN buffer, 0.1 % Tween-20) to reduce the DTT concentration to <0.5 mM. Lysates were incubated with Anti-FLAG®-M2 Magnetic Beads (Merck, M8823) rotating at 4 °C for 2 hours. Beads were washed with IP buffer, and protein was eluted in 100 µg/ml 3x FLAG™ Peptide (Merck, F4799) rotating at 4 °C for 1 hour.

### *E. coli* protein expression and purification

The pET-15b expression vector encoding protein of interest was transformed into *E. coli* BL21 (DE3) cells. Overnight culture was inoculated into 0.1 or 1.0 L Luria-Bertani (LB) medium in a 0.25 or 2 L Erlenmeyer flask, and cell culture was grown at 37 °C until OD_600_ 0.4–0.8 was reached. The culture temperature was reduced to 25 °C, and protein expression induced via addition of isopropyl β-D-1-thiogalactopyranoside (IPTG) (final concentration 0.1 mM). After incubation at 25 °C for 4 hours, cells were harvested via centrifugation at 3500 g for 5 min, resuspended in 25 mM HEPES, 150 mM NaCl, pH 7.5 (“buffer”) and stored at −20 °C. Thawed cell suspensions were lysed via sonication and subjected to centrifugation at 20 000 g for 30 min, and supernatant passed through a 0.22 µm syringe filter. Proteins were purified using an ÄKTA Go protein purification system (Cytiva) using HisTrap FF column (Cytiva, 17525501) for Ni^2+^ immobilized metal ion affinity chromatography (IMAC) with buffer + 25 mM imidazole, or Strep-Tactin®XT 4Flow® high capacity FPLC column (IBA, 2-5027-001) for strep tag affinity chromatography (strepAC), following the manufacturer’s instructions. For small scale purification, HisPur™ Ni-NTA Resin (Thermo Fisher Scientific, 10449164) was used for IMAC and Strep-Tactin®XT 4Flow® high-capacity Spin Column Kit (IBA, 2-5151-000) was used for strepAC, following manufacturer’s instructions. Homemade TEV or 3C protease was used to elute proteins of interest from affinity resin or used to cleave tags after proteins were eluted from resin. Size exclusion chromatography (SEC) was performed using a Superose 6 Increase 10/300 GL column (Cytiva, 29091596), with 0.5 mL injection volume and 0.5 mL/min flow rate following manufacturer’s instructions. SEC was calibrated using Protein Standard Mix 15 - 600 kDa (Merck, 69385). SEC data was processed using Microsoft Excel or a custom Streamlit python app (supplementary information).

Size exclusion chromatography with multi angle light scattering (SEC-MALS) SEC-MALS experiments were performed at ambient temperature, using an HPLC+LS+RI+QELS configuration pre-equilibrated with buffer. The instrument set up included the HPLC modules of Agilent 1260 Infinity II series (Agilent Technologies) connected in-line to the miniDAWN (NEON) with a three-angle (49°, 90° and 131°) light scattering detector, and an integrated QELS dynamic light scattering detector at 135° for determination of hydrodynamic radius. Concentration was measured using UV absorbance at 280 nm using the in-line 1260 infinity II series DAD detector (Agilent Technologies) and the Wyatt Optilab™ (NEON) refractive Index detector (Wyatt Technology, Santa Barbara, CA). Approximately 10 µg of protein was injected onto the SEC column (Waters XBridge Premier Protein SEC 250Å, 2.5 µm, 4.6 x 150 mm) at a flow rate of 0.2 mL/min for run time of 20 or 25 minutes per sample. Data collection for QELS was set to DLS interval of 2 sec and Collection Interval of 0.5 sec. The DAD module G7115A (Agilent Technologies) of HPLC was set to collect UV absorbance signals at 280 nm (Aromatic Trp, Tyr absorbance), 210 and 214nm (non-aromatic absorbance, amide bond). Sample submission, data collection and SEC-MALS analysis were performed using the HPLC connect and ASTRA 8.2.2 software (Wyatt Technology). Alignment of ultraviolet (UV), refractive index (RI) and light scattering (LS) peaks, band broadening and RI-based mass calculation were performed before export of raw data. A custom Streamlit python app was used to visualize data and calculate a single mass value for each protein replicate. If the largest UV peak in the chromatogram was visibly irregular and asymmetrical, an exponentially modified Gaussian (EMG) was fit so that, to reduce the contribution of shoulder regions including other protein species to subsequent calculations. The average mass in the x value range corresponding to the UV or EMG peak apex ∓ 10 % of the peak’s full width at half maximum was taken forward. The mass of a protein monomer was estimated based on the expressed sequence including known protease cleavage sites, and SDS-PAGE mobility relative to markers of known mass.

### Crystallisation, X-ray data collection and processing

To identify the lead crystallisation condition, the protein complex was subjected to a range of commercially available screens in a 96-well sitting drop plate format. The first lead crystallisation condition was identified in the Classics screen (Qiagen) following a two-day incubation at 20 °C in a drop containing 533 nL of protein complex in buffer at 1.5 mg ml⁻¹ and 267 nL of the reservoir solution containing 0.2 M lithium sulfate, 0.1 M HEPES pH 7.5, 25 % (w/v) PEG 3350. Crystals were passed through a solution of mother liquor supplemented with 20% PEG 400 as a cryoprotectant, then immersed and stored in liquid nitrogen. Data were collected at 100 K on beamline I04 at the Diamond Light Source (Didcot, UK), using a wavelength of 0.9537 Å and an Eiger2 XE 16M detector. Diffraction data were integrated using XDS through the automated data-processing pipeline at Diamond Light Source and were subsequently scaled and merged in AIMLESS (50).

### Structure determination and refinement

Molecular replacement was performed using PhaserMR (51) using the AlphaFold 3 prediction of Rp58 residues 3-83 (tetrameric) as the search model. A random 5% of reflections were set aside prior to refinement to calculate R_free_. Iterative rounds of manual model building and density inspection in COOT (52) (were combined with refinement in REFMAC5 (53). B-factors were refined isotropically and hydrogen atoms were included in riding positions. Water molecules were assigned to well-defined peaks in the difference electron density map (>3.5 σ) forming plausible hydrogen bonds (2.5–3.5 Å) with donor or acceptor groups. Sodium ions and alternative rotamers for flexible side chains were modelled where density clearly indicated. Model geometry was monitored throughout using MolProbity (54) and built-in COOT validation tools. The coordinate and structure factor files were processed through the PDB-REDO web server (23) for final map optimization and automated re-refinement. Outlier residues and geometrical clashes flagged by the wwPDB validation server were manually adjusted in COOT, and atomic coordinates and structure factors were deposited in the PDB under accession code 34DI. A representative portion of the electron density map is shown in **Figure S1**.

### Nano differential scanning fluorimetry (NanoDSF)

Thermal stability was evaluated using a Prometheus Panta system (NanoTemper Technologies). Samples were prepared at 0.5-5 mg/mL in buffer and loaded into Tycho NT.6 capillaries (NanoTemper Technologies, TY-C001). Thermal melting curves were recorded from 25-90 °C at a heating rate of 1.0 °C/min. Intrinsic fluorescence was recorded at 330 nm and 350 nm to monitor tertiary structure changes. Data acquisition and export were executed using Panta Control 1.7.1 software. Data visualization was performed in Microsoft Excel.

## Supporting information

Crystal structure data

## Supplementary files

Supplementary Figures and Tables Supplementary Information Crystal Structure Data

## Data availability

The code includes Streamlit apps for SEC and SEC-MALS that enable easier exploration of the data.

## Author contributions

Author contributions: T.W., J.R.B., J.I.B.B. and W.N.H. conceived and designed the research; T.W. and J.R.B. performed experiments; T.W. and S.R.F. performed *in silico* analyses; S.R.F designed and wrote the scripts for each covariation analysis; T.W., J.R.B., S.R.F., G.P., J.I.B.B. and W.N.H. contributed to data interpretation; T.W., J.R.B., S.R.F. and J.I.B.B. wrote the manuscript. All authors commented on the manuscript.

## Acknowledgements

This work was supported by the European Union (ERC Consolidator grant, project number 101000997, APHIDTRAP, awarded to JIBB). The authors acknowledge Research Computing at the James Hutton Institute for providing computational resources and technical support for the “UK’s Crop Diversity Bioinformatics HPC” (BBSRC grants BB/S019669/1 and BB/X019683/1), use of which has contributed to the results reported within this paper. We thank Ms. Yao Fu for generating the *E.coli* expression vectors for Rp1 and Rp58. We acknowledge Thomas Eadsforth and Marilyn Paul at the University of Dundee Drug Discovery Unit for assistance in SEC-MALS and crystallography data collection, and Heather Grey and Professor Steven Spoel for providing the pEarleyGate 202 plasmid.

## Competing Interests

The authors have declared no competing interests.

