## Supplementary Figures and Tables for "An aphid effector pair forms a hetero-oligomeric complex important for protein stability and activity"

**A**

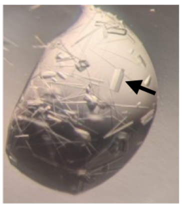

**B**

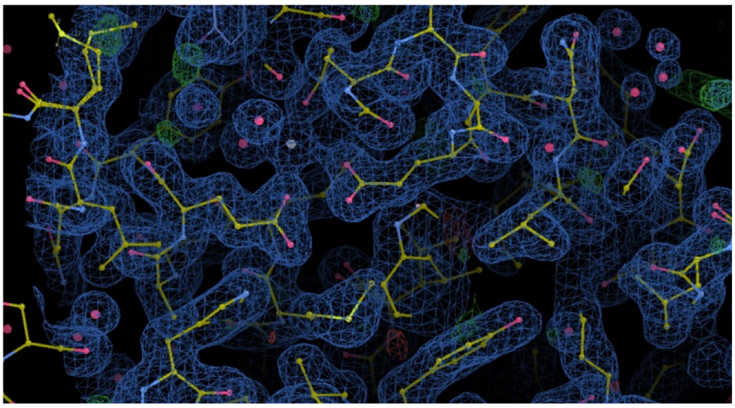

**Figure S1. Crystallography results. (A)** crystallization drop - crystal used to determine the structure indicated with an arrow. **(B)** sample of post-refinement electron density and protein model, visualized using COOT. Electron density (blue, with $F_O$-$F_C$ difference map green) is contoured at 1.0 σ and atoms are coloured: oxygen (red), carbon (brown/yellow), nitrogen (blue), sulfur (yellow), sodium (silver).

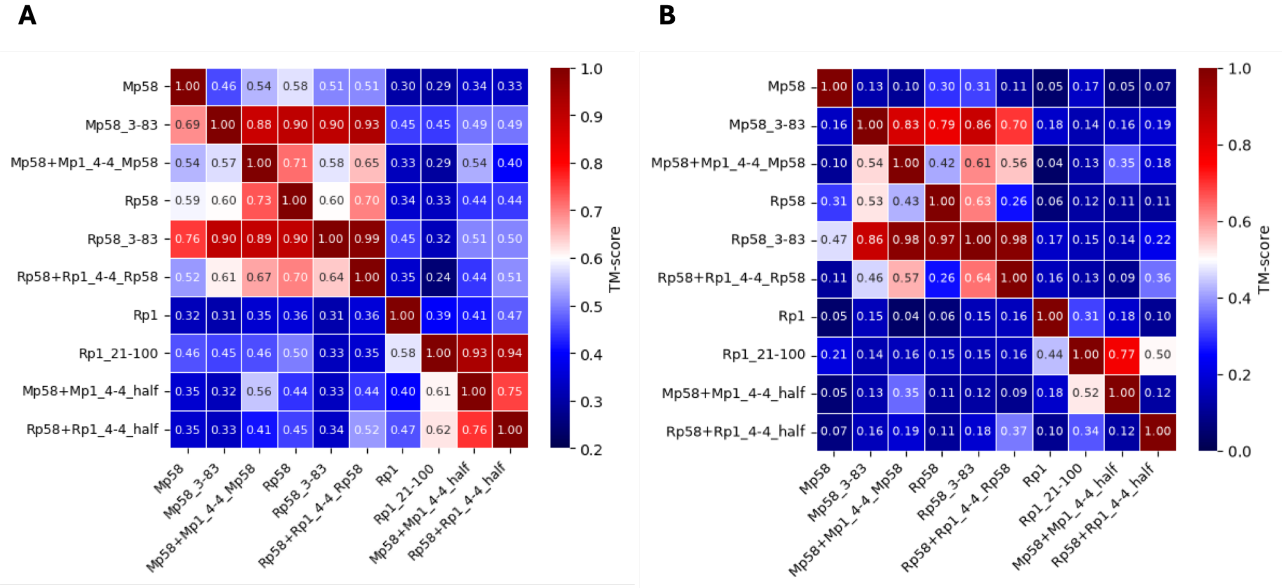


**Figure S2.** Tetrameric AlphaFold3 models of full-length (no suffix) or truncated (*e.g.* residue indices 3-83) proteins were used. Octameric effector complex models (Mp58+Mp1_4-4, Rp58+Rp1_4-4) were prepared for analysis by extracting the middle tetramer (Mp58 or Rp58) or one symmetrical half of the complex containing 2 subunits of each of the 2 interacting proteins. The displayed TM scores are calculated based on the proteins in the rows, not the columns. **(A)** TM score from MMalign. **(B)** TM score calculated using the standard formula after superimposition using the PyMOL “super” command.

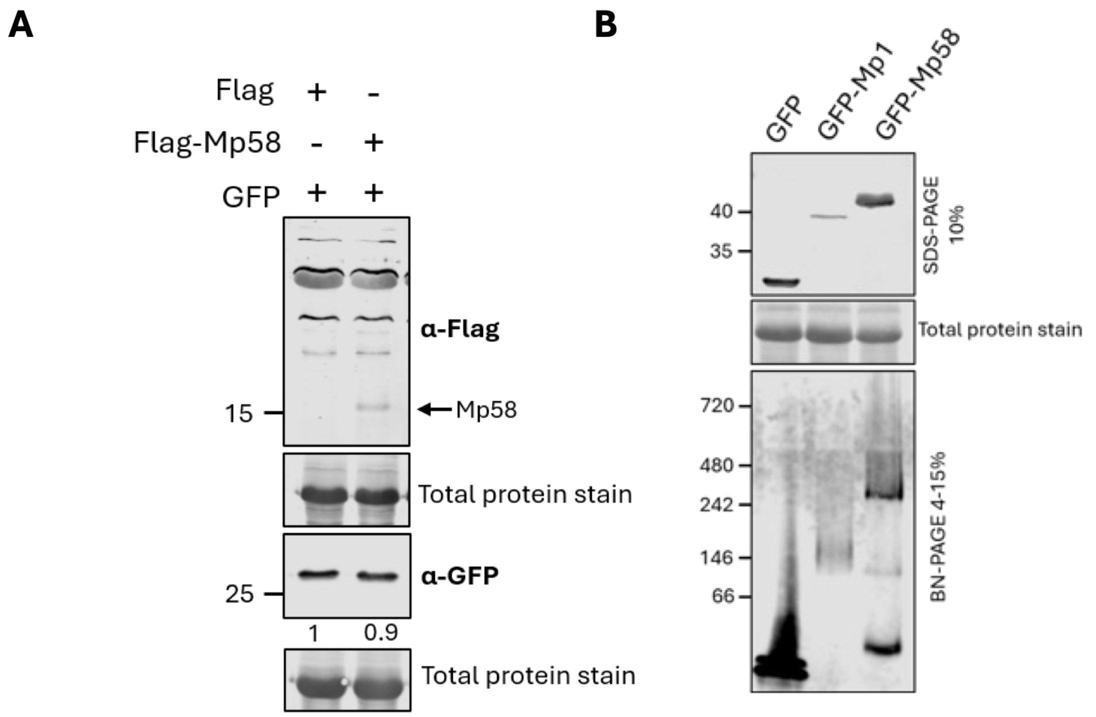


**Figure S3. (A)** IB using FLAG or GFP antibodies after sodium dodecyl sulfate polyacrylamide gel electrophoresis (SDS-PAGE) of protein extracts of *N. benthamiana* co-expressing GFP with FLAG or FLAG-Mp58 proteins. Protein quantification was analyzed by normalizing the band intensity of GFP to the total protein amounts using Empiria Studio. To generate band intensity ratios, GFP levels were compared between samples in which FLAG-Mp58 or FLAG (control, set at 1) were co-expressed. **(B)** IB using GFP antibody after SDS-PAGE or blue native PAGE (BN-PAGE) of protein extracts of *N. benthamiana* expressing the indicated single proteins. Numbers refer to mass of PAGE marker proteins (kilodaltons, left of image) or band intensity ratio (below GFP blot image). Data are representative of n = 2 independent biological replicates.

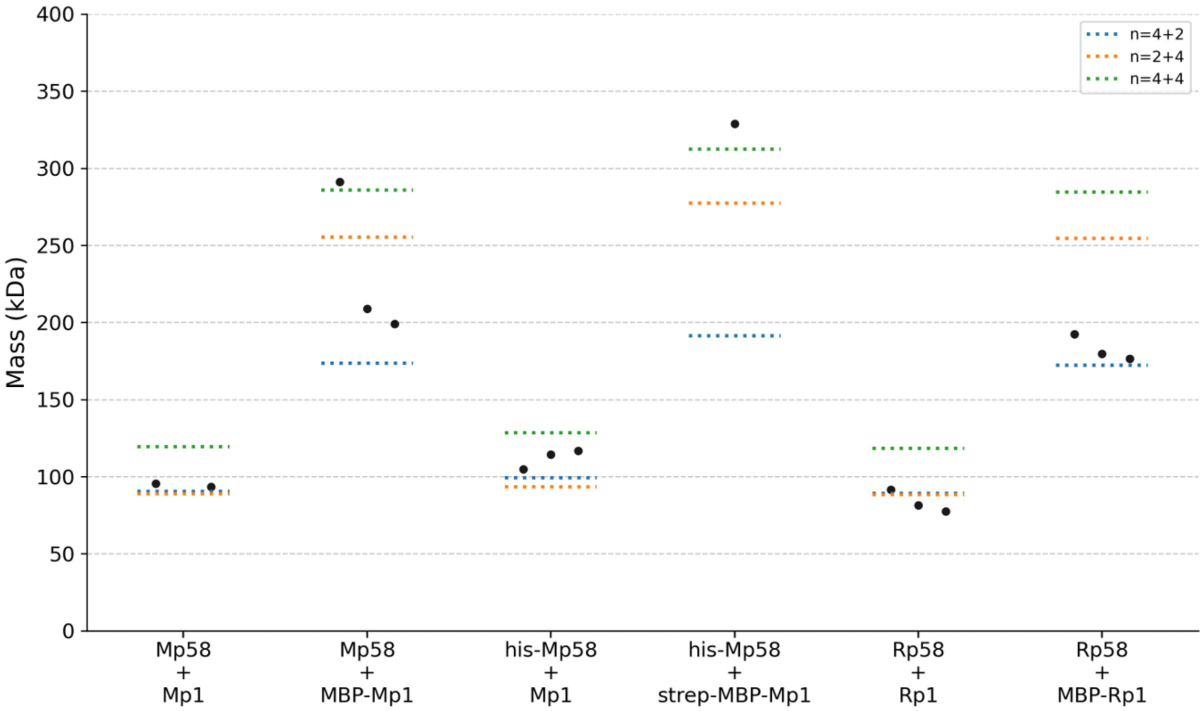


**Figure S4. Oligomerization state estimation of effector complexes expressed in and purified from *E. coli*.** Size exclusion chromatography with multi-angle light scattering (SEC-MALS) results for *Myzus persicae* (Mp58+Mp1) and *Rhopalosiphum padi* (Rp58+Rp1) effector complexes, with various protein tags present including his, strep or maltose binding protein (MBP). Graph shows mass measured (1 biological replicate per black dot), expected mass of two-protein complex oligomerization state n (color-coded dashed lines).

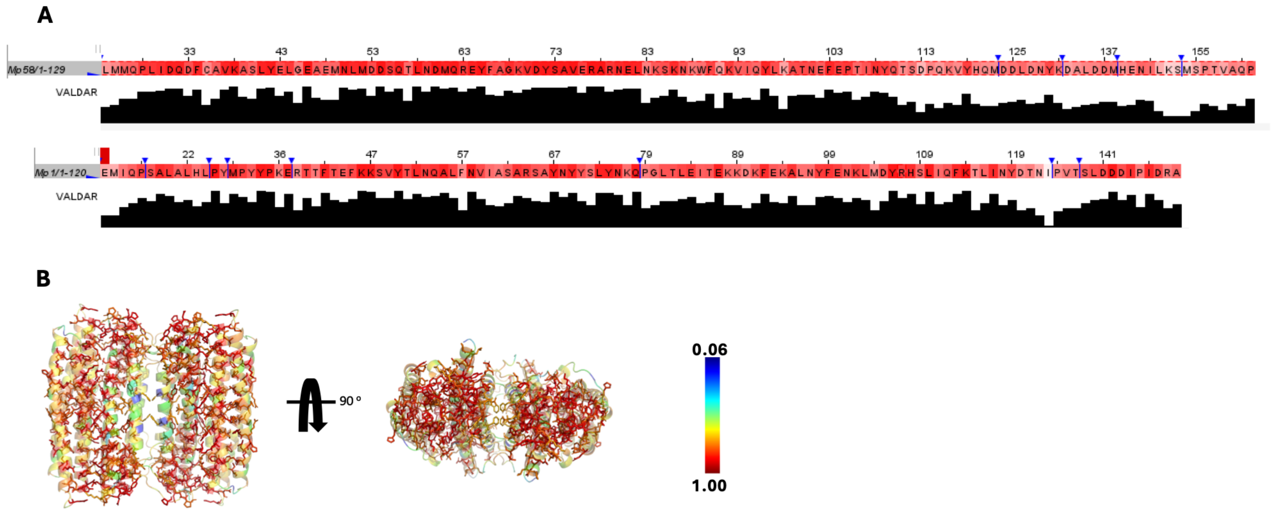


**Figure S5. Residue conservation in the effector complex. (A)** sequence alignments of orthologues of Mp58 (above) and Mp1 (below), colored by Valdar amino acid conservation score. Column chart of normalized Valdar score (0-1) is shown below the alignment. Calculation and visualization was performed in Jalview, other sequences and insertions were hidden. **(B)** octameric AlphaFold 3 prediction of the *Myzus persicae* effector complex with 4 Mp58 and 4 Mp1 subunits. Models are shown in transparent PyMOL cartoon style, colored by Valdar amino acid conservation score (color bar), and residues with score >0.8 are shown in PyMOL stick style.

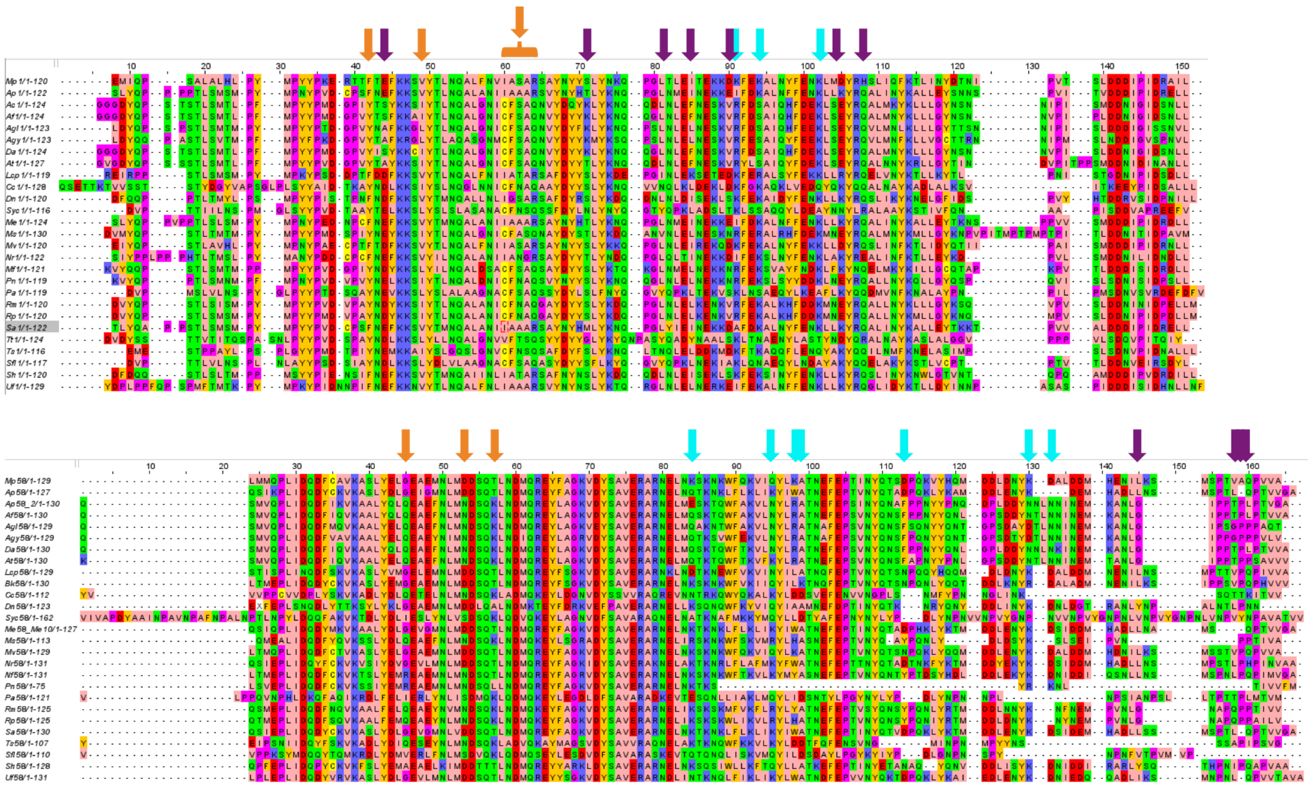

**Figure S6. Covariation analysis of aligned Mp58 and Mp1 protein sequences.** Sequence alignments of orthologues of Mp1 (above) and Mp58 (below), coloured by amino acid side chain chemical property (Zappo scheme). Markers above cluster-associated residues are coloured according to the cluster colour code in **Figure 4B.**

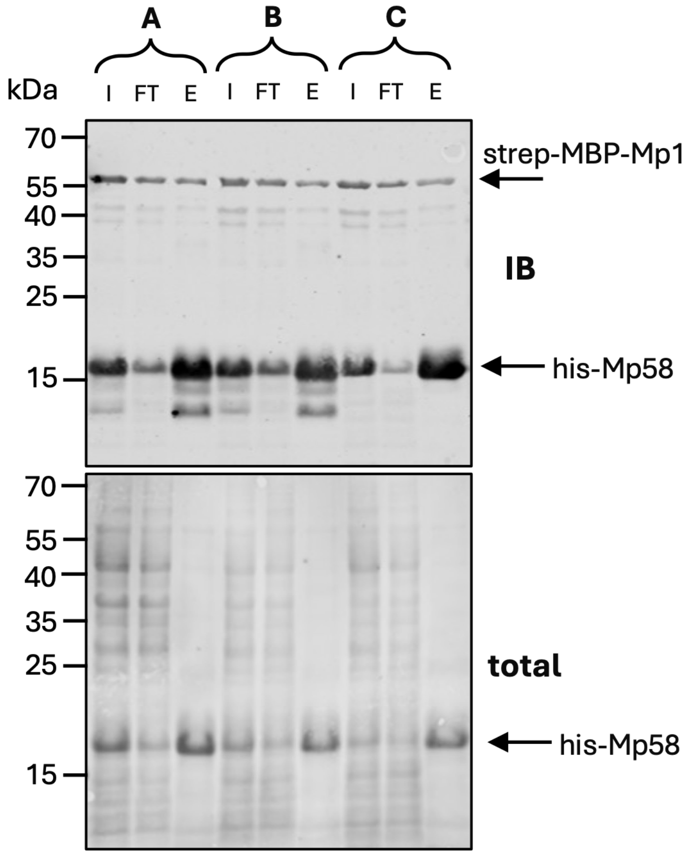


**Figure S7. *In vitro* interaction between Mp58 and Mp1 mutants.** *E. coli* lysates containing single proteins were mixed, his-tagged Mp58 proteins were purified and co-purification of interacting Mp1 proteins containing strep and maltose binding protein (MBP) tags was assessed. The following combination were tested: Mp58+Mp1 (**A**), Mp58+Mp1$^{F40A}$ (**B**), Mp58$^{Y19A.D30A}$+Mp1$^{F40A}$ (**C**). Input (I), flow-through (FT) and elution (E) fractions were used. Immunoblotting (IB) using his and strep antibodies was performed after sodium dodecyl sulfate polyacrylamide gel electrophoresis (SDS-PAGE) and total protein staining. Numbers refer to mass of SDS-PAGE marker proteins (kilodaltons). Data are representative of n = 2 independent biological replicates.

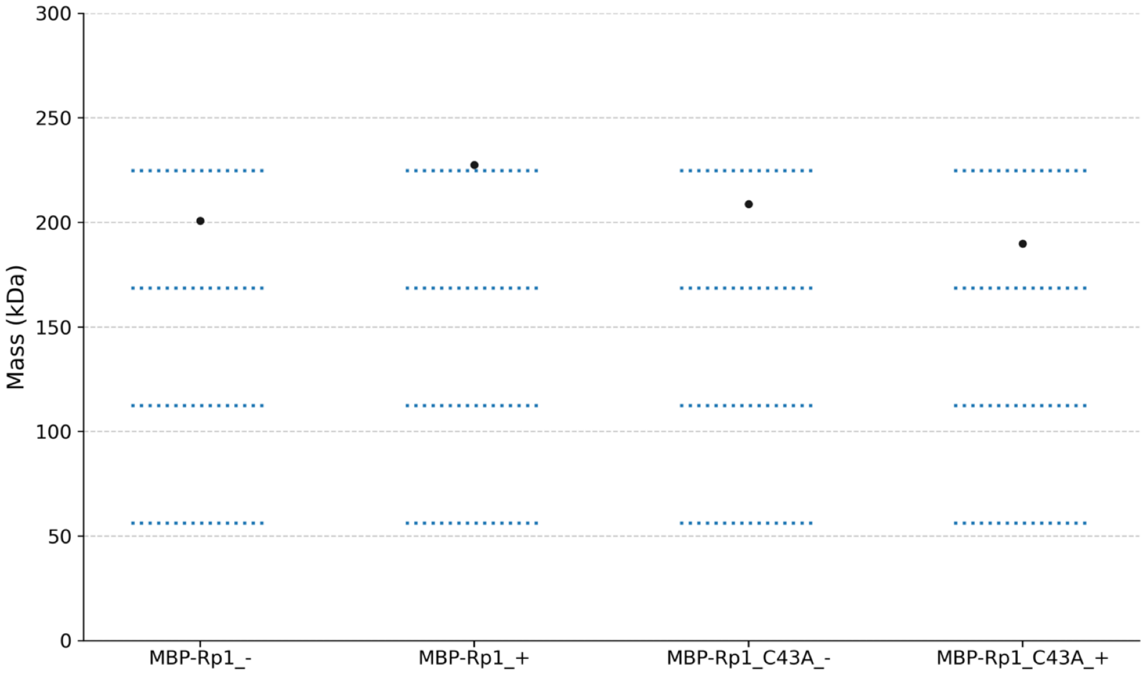


**Figure S8. Effect of cysteine disruption on Rp1 oligomerization state.** Size exclusion chromatography with multi-angle light scattering (SEC-MALS) results for maltose binding protein (MBP)-tagged Rp1 proteins, expressed in and purified from *E. coli*. Rp1 wild-type or C43A mutant was used, and proteins were incubated in 0 (-) or 10 mM (+) Tris(2-carboxyethyl)phosphine overnight at room temperature. Graph shows mass measured for 1 biological replicate (black dot), expected mass of oligomerization states 1-4 (blue dashed lines).

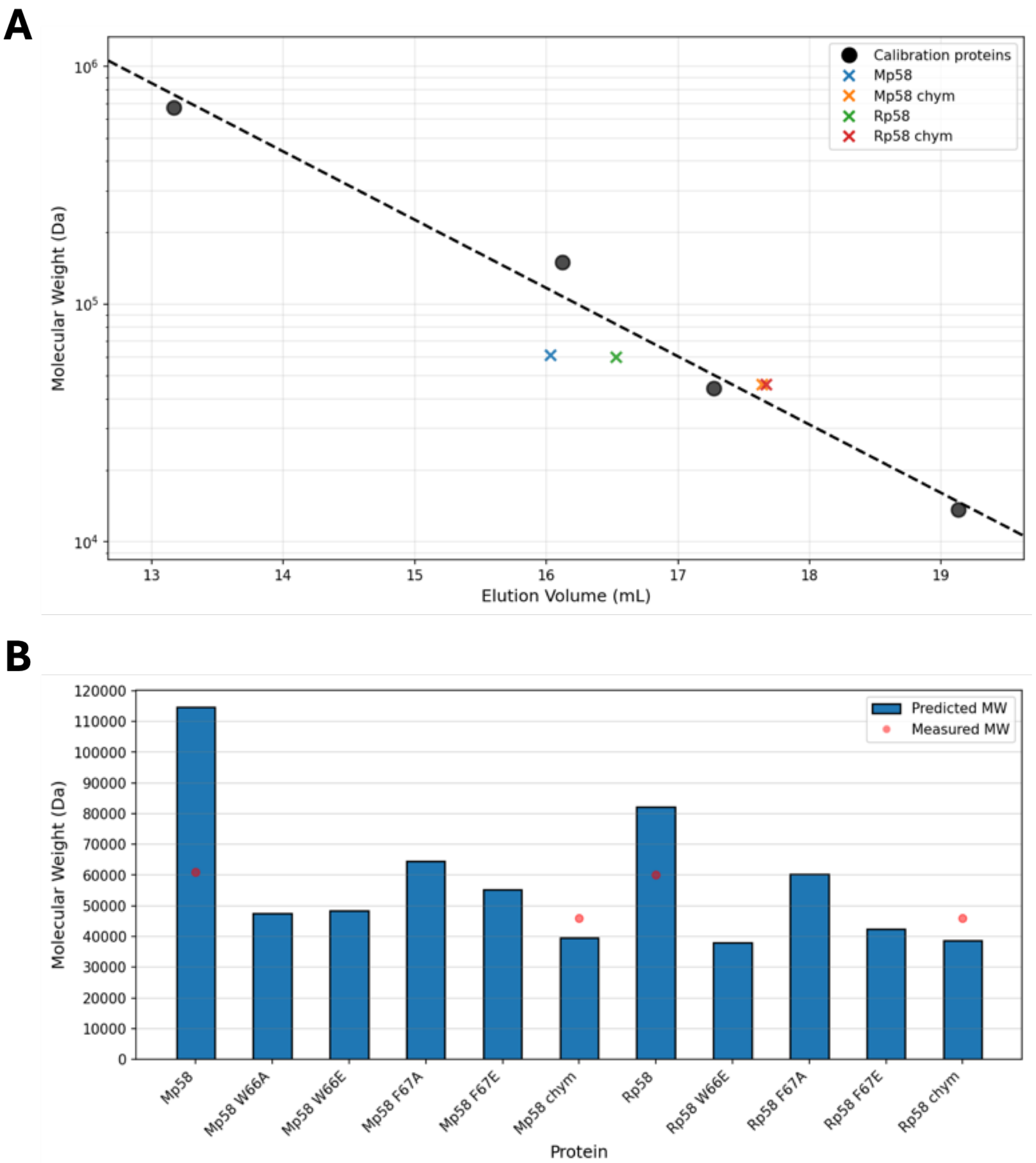


**Figure S9. Size exclusion chromatography (SEC) calibration and protein mass estimation. (A)** SEC calibration scatter graph showing elution volume of the largest ultraviolet peak of standard calibration proteins and test proteins of known mass. Graph with logarithmic y axis and least squares regression line based on calibration proteins. The test proteins are Mp58 and Rp58, expressed in and purified from *E. coli*, digested with tobacco etch virus protease (no suffix, containing flexible regions) or chymotrypsin ("chym" suffix, lacking flexible regions). **(B)** Column chart showing estimated mass of Mp58 and Rp58 variants based on regression line in A, and measured mass of proteins where available. Data are representative of 2≤n≤3 independent biological replicates.

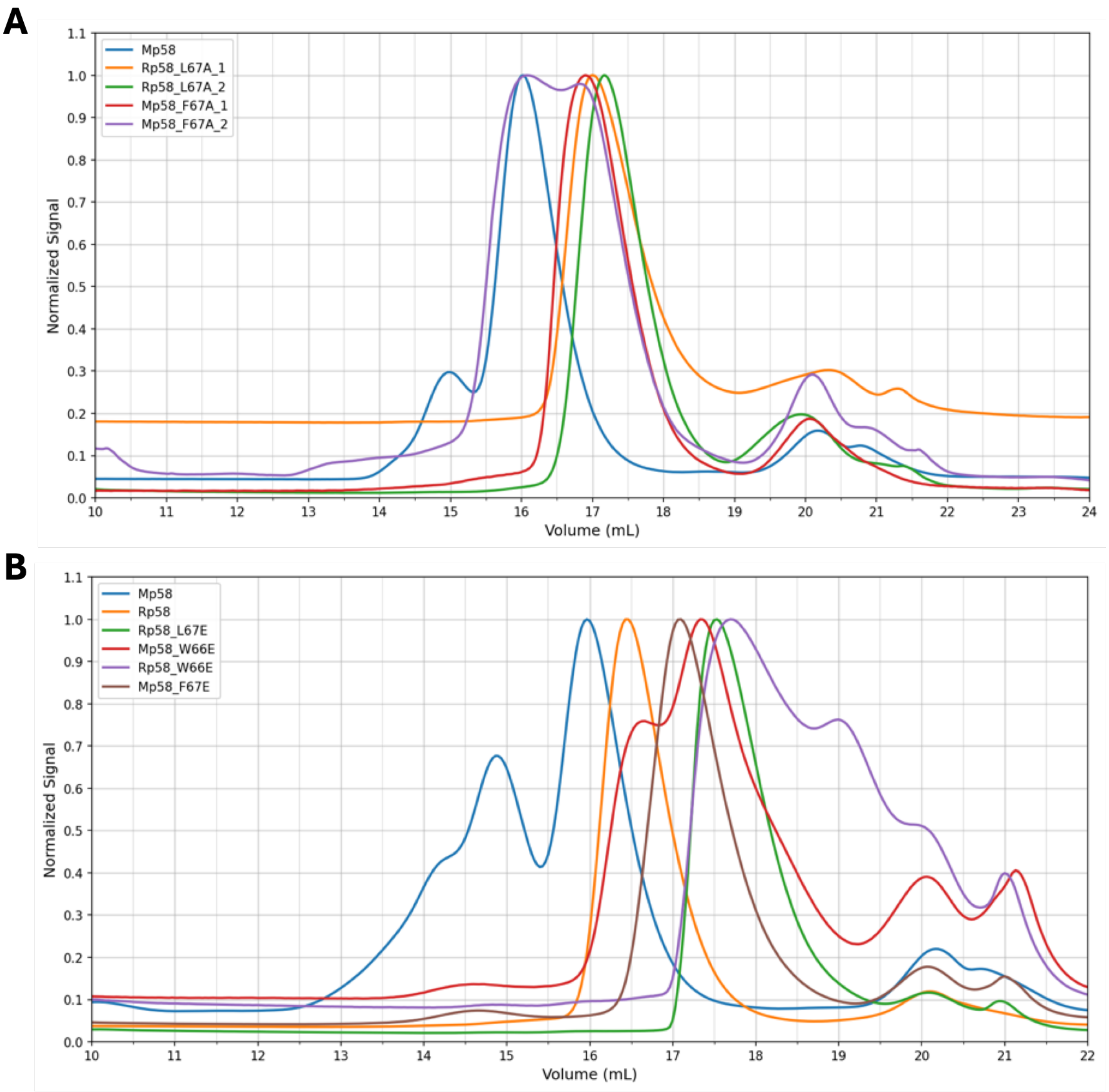


**Figure S10. size exclusion chromatography of Mp58 and Rp58.** Size exclusion ultraviolet chromatograms of Mp58 and Rp58 sequence variants. **(A)** Mp58 and Rp58 residue index 67 alanine substitution mutants, immediately after tobacco etch virus protease digest (1) and after 1 week of storage in fridge (2). **(B)** Mp58 and Rp58 wild-type or glutamate substitution mutants. Data are representative of 2≤n≤3 independent biological replicates.

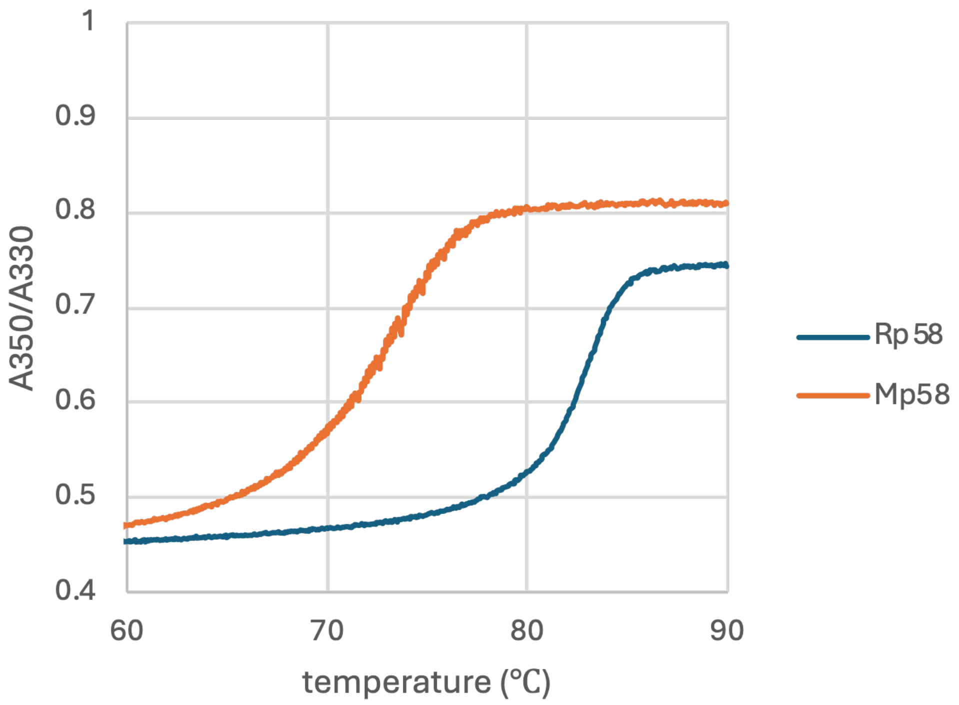


**Figure S11. Mp58/Rp58 NanoDSF.** Thermal unfolding graphs for Mp58 and Rp58 proteins, expressed in and purified from *E. coli*. Data are representative of n = 3 independent biological replicates.

**Table S1. AlphaFold3 model statistics**

| protein 1 | protein 2 | copies | pTM (%) | ipTM (%) | average core pLDDT |
|---|---|---|---|---|---|
| Mp58 | - | 4 | 48 | 53 | 82 |
| $Mp58_{3-83}$ | - | 4 | 57 | 66 | 90 |
| Rp58 | - | 4 | 55 | 58 | 87 |
| $Rp58_{3-83}$ | - | 4 | 88 | 89 | 95 |
| Mp1 | - | 4 | 37 | 45 | 85 |
| $Mp1_{21-100}$ | | 4 | 76 | 80 | 91 |
| Rp1 | - | 4 | 45 | 53 | 89 |
| $Rp1_{21-100}$ | - | 4 | 92 | 92 | 98 |
| Mp58 | Mp1 | 4 + 4 | 66 | 67 | 86, 89 |
| Rp58 | Rp1 | 4 + 4 | 72 | 72 | 92, 95 |

#### Supplementary SEC-MALS data

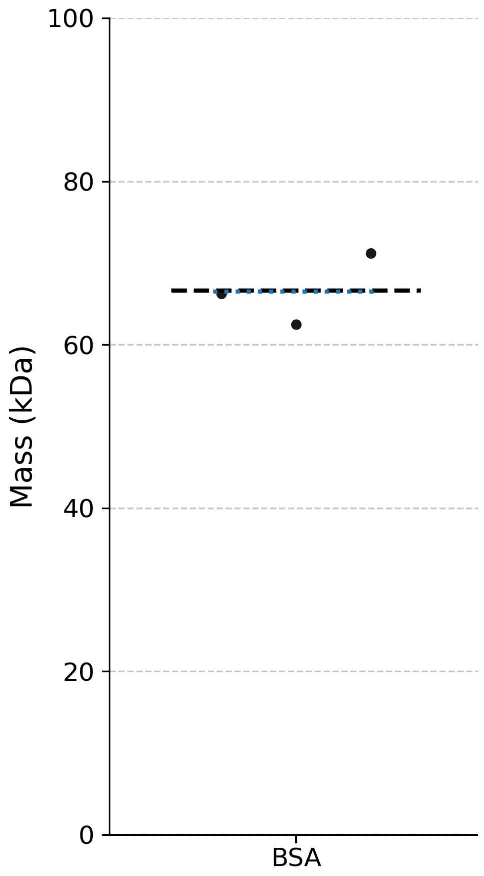


**Figure showing SEC-MALS BSA control.** Size exclusion chromatography with multi-angle light scattering (SEC-MALS) results for bovine serum albumin (BSA). Graph shows mass measured for 3 biological replicates (black dots), mean (black dashed line), expected mass of monomer (blue dashed line).

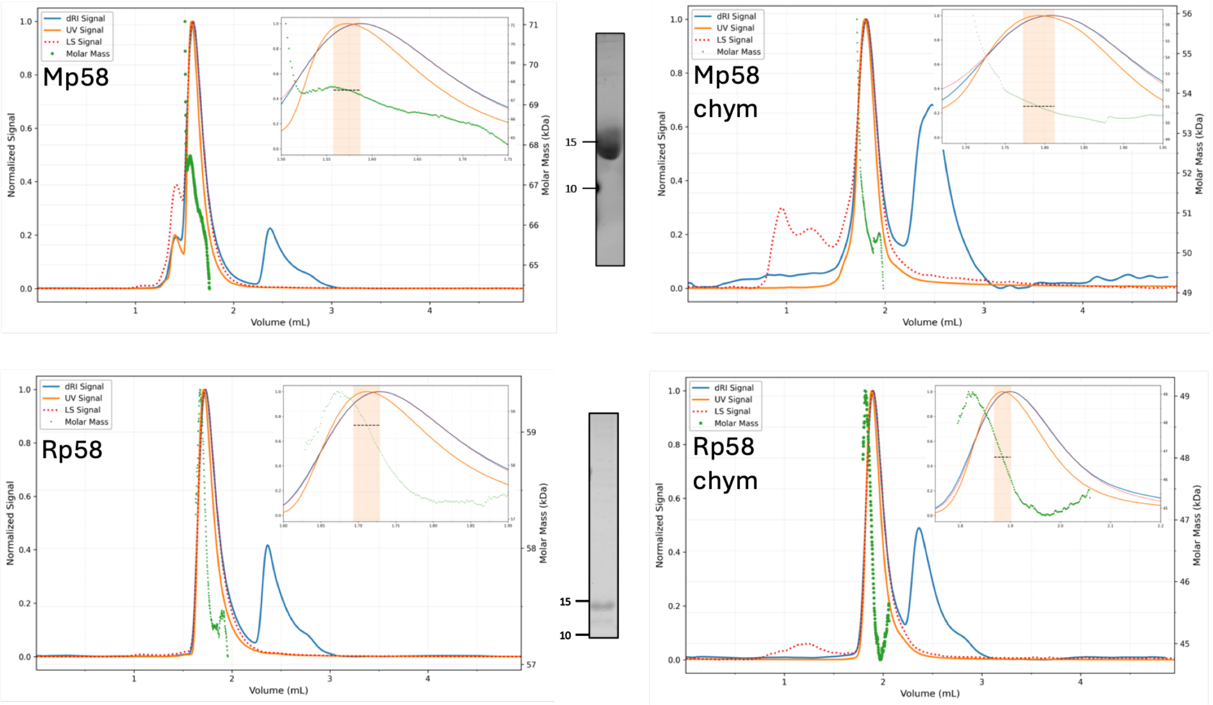


**Figure showing the protein mass calculation using size exclusion chromatography with multi-angle light scattering.** For each protein, chromatograms including refractive index (dRI), ultraviolet (UV) and light scattering (LS) are shown. If the largest UV peak has an irregular shape, an exponentially modified Gaussian (EMG) fit of this region is performed within manually specified limits (grey shade); the inset shows a magnified view of this region. Molar mass (green dots) is calculated using ASTRA software (Wyatt Technology) and the average mass (black dashed line) is found in the x value range corresponding to the UV or EMG peak apex ∓ 10 % of the peak's full width at half maximum (orange shade). For each protein, representative results of total protein staining after purification and sodium dodecyl sulfate polyacrylamide gel electrophoresis are shown to the right of graphs; numbers refer to mass of marker proteins. Proteins were expressed in and purified from *E. coli*, digested with tobacco etch virus protease (no suffix) or chymotrypsin (" chym" suffix). Data are representative of n = 3 independent biological replicates.

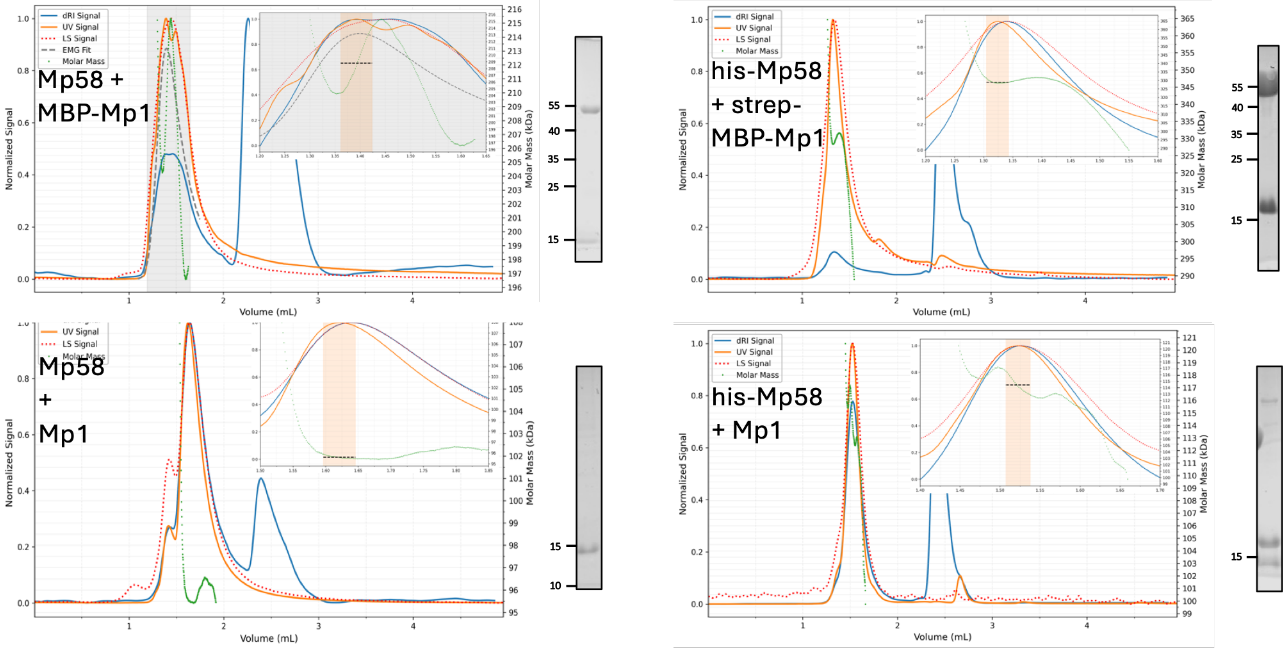


**Figure showing protein mass calculation using size exclusion chromatography with multi-angle light scattering.** For each protein, chromatograms including refractive index (dRI), ultraviolet (UV) and light scattering (LS) are shown. If the largest UV peak has an irregular shape, an exponentially modified Gaussian (EMG) fit of this region is performed within manually specified limits (grey shade); the inset shows a magnified view of this region. Molar mass (green dots) is calculated using ASTRA software (Wyatt Technology) and the average mass (black dashed line) is found in the x value range corresponding to the UV or EMG peak apex ∓ 10 % of the peak's full width at half maximum (orange shade). For each protein, representative results of total protein staining after purification and sodium dodecyl sulfate polyacrylamide gel electrophoresis are shown to the right of graphs; numbers refer to mass of marker proteins. Proteins were expressed in and purified from *E. coli* with tags including his, strep or maltose binding protein (MBP); purified proteins were digested with tobacco etch virus protease and/or Human Rhinovirus 3C protease. Data are representative of n = 3 independent biological replicates.

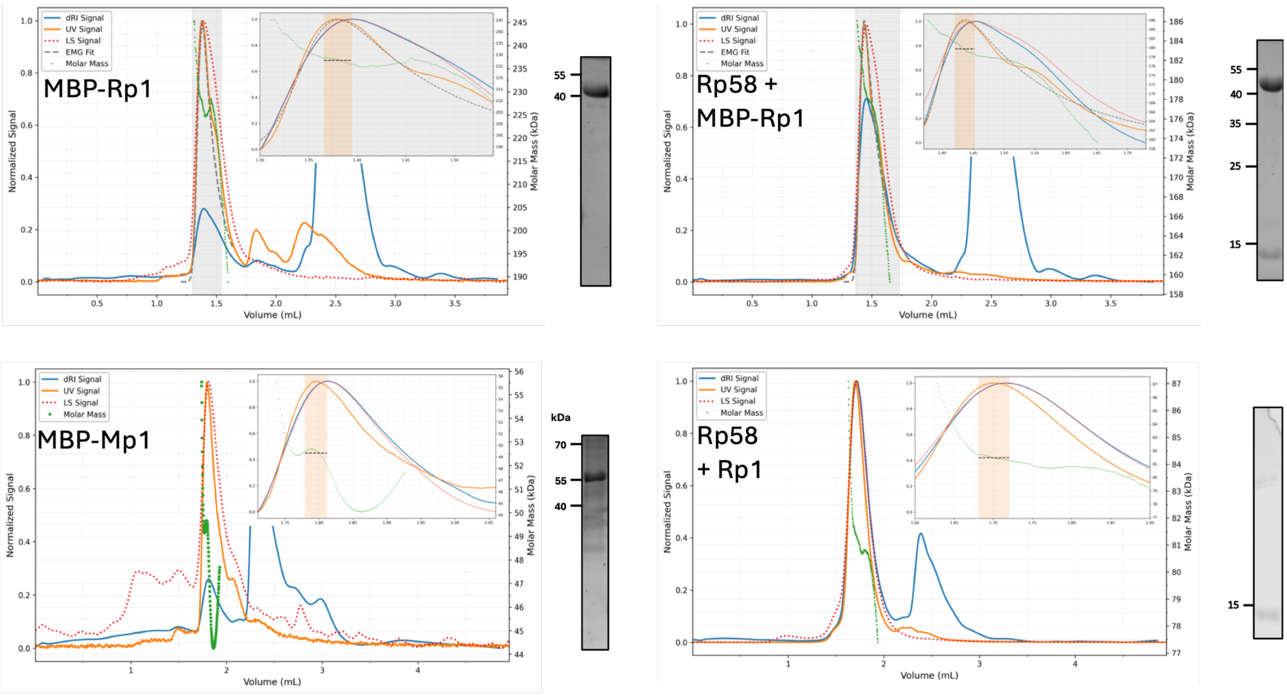


**Figure showing protein mass calculation using size exclusion chromatography with multi-angle light scattering.** For each protein, chromatograms including refractive index (dRI), ultraviolet (UV) and light scattering (LS) are shown. If the largest UV peak has an irregular shape, an exponentially modified Gaussian (EMG) fit of this region is performed within manually specified limits (grey shade); the inset shows a magnified view of this region. Molar mass (green dots) is calculated using ASTRA software (Wyatt Technology) and the average mass (black dashed line) is found in the x value range corresponding to the UV or EMG peak apex ∓ 10 % of the peak's full width at half maximum (orange shade). For each protein, representative results of total protein staining after purification and sodium dodecyl sulfate polyacrylamide gel electrophoresis are shown to the right of graphs; numbers refer to mass of marker proteins. Proteins were expressed in and purified from *E. coli* with tags including maltose binding protein (MBP); purified proteins were digested with tobacco etch virus protease and/or Human Rhinovirus 3C protease. Data are representative of n = 3 independent biological replicates.

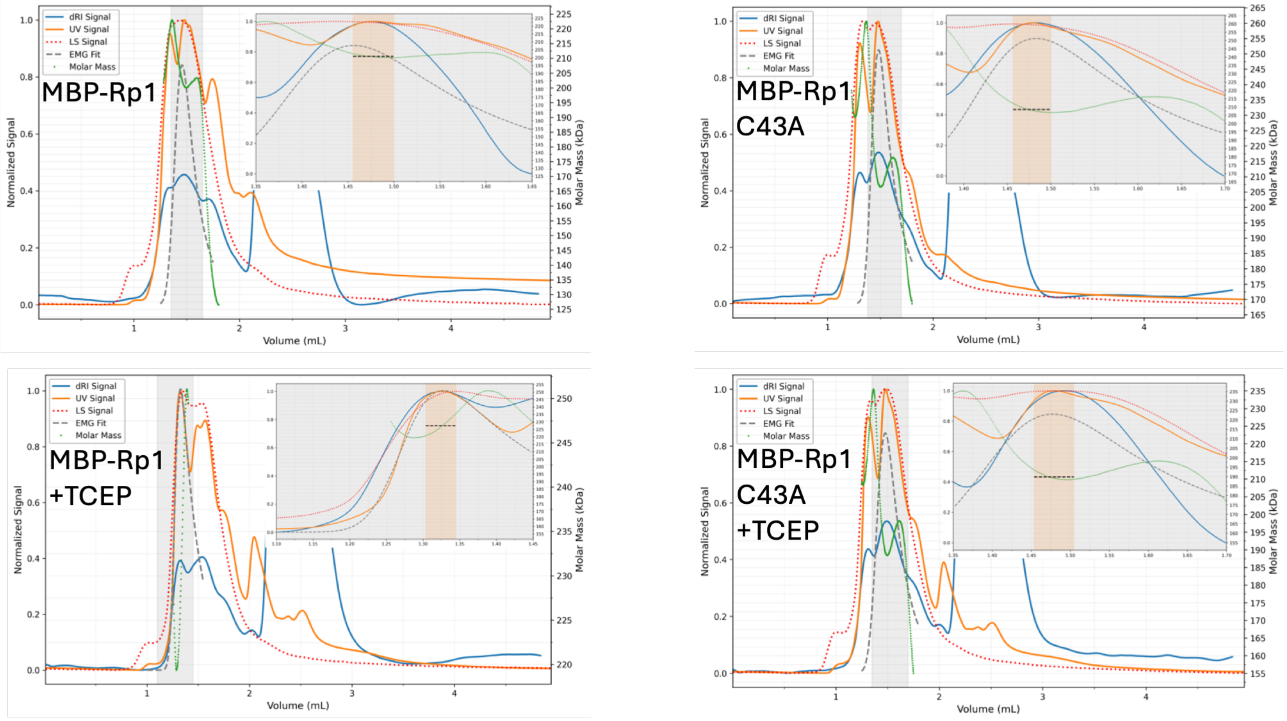


**Figure showing protein mass calculation using size exclusion chromatography with multi-angle light scattering.** For each protein, chromatograms including refractive index (dRI), ultraviolet (UV) and light scattering (LS) are shown. If the largest UV peak has an irregular shape, an exponentially modified Gaussian (EMG) fit of this region is performed within manually specified limits (grey shade); the inset shows a magnified view of this region. Molar mass (green dots) is calculated using ASTRA software (Wyatt Technology) and the average mass (black dashed line) is found in the x value range corresponding to the UV or EMG peak apex $\mp$ 10 % of the peak's full width at half maximum (orange shade). Rp1 wild-type or C43A mutant were expressed in and purified from *E. coli* with tags including maltose binding protein (MBP); purified proteins were digested with tobacco etch virus protease in the presence of 0 or 10 mM Tris(2-carboxyethyl)phosphine (TCEP).

### Uncropped Western Blots, and additional replicates, corresponding to figures

Fig. 3A

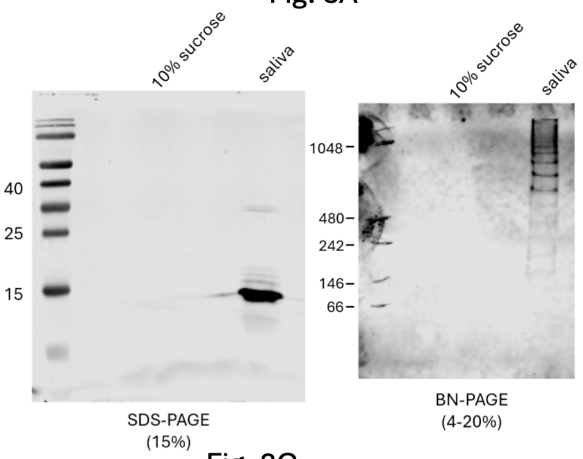


Fig. 3B

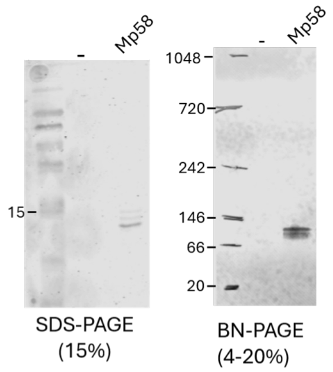


Fig. 3C

FLAG-Mp58 x GFP-Mp1 co-IP

Rep1

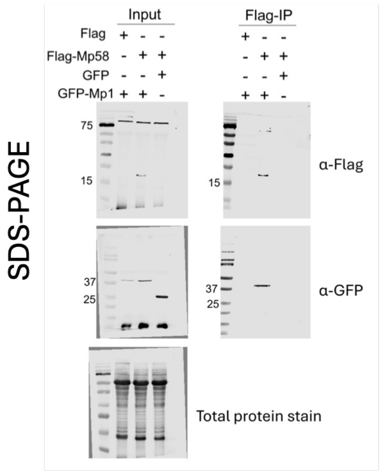


Fig. 3D

Rep2 (Fig.3C)

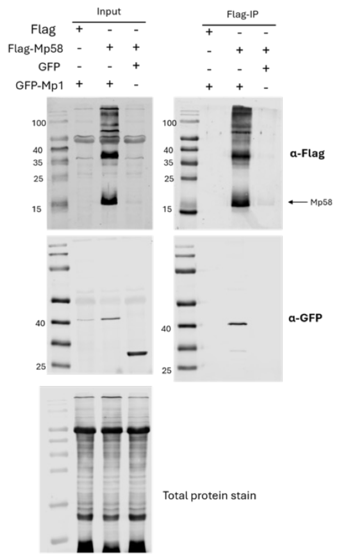


Rep1

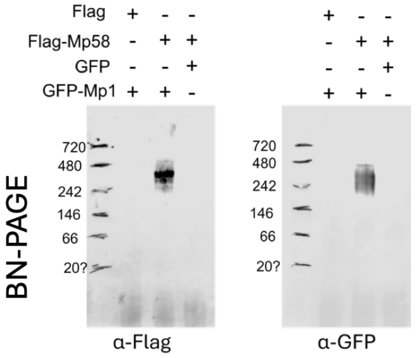


Rep2 (Fig. 3E)

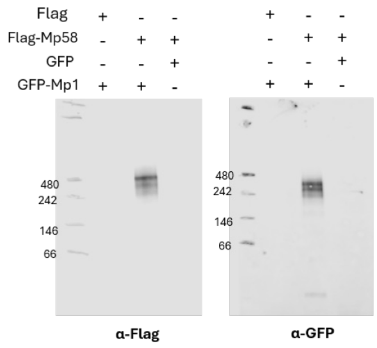

### FLAG-Mp1 x GFP-Mp58 co-IP

#### Rep1

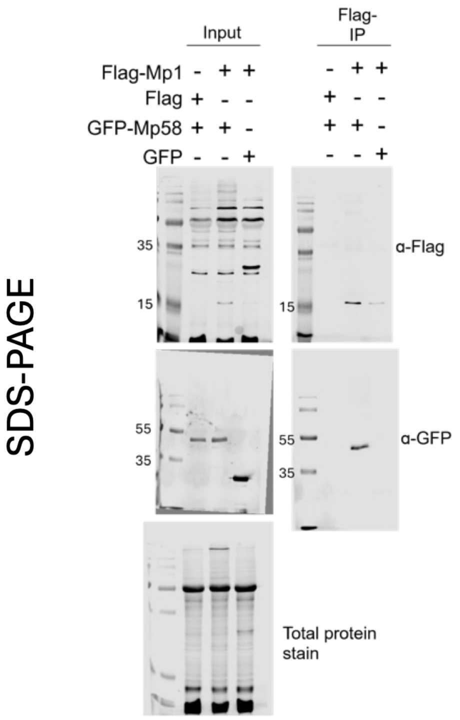


#### Rep2 (Fig.3D)

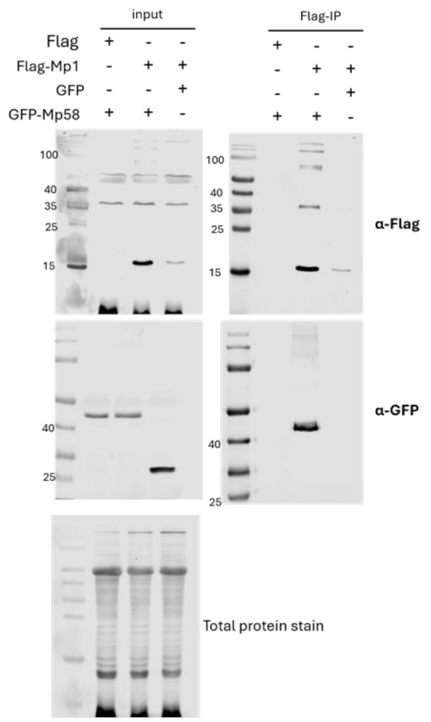


#### Rep1

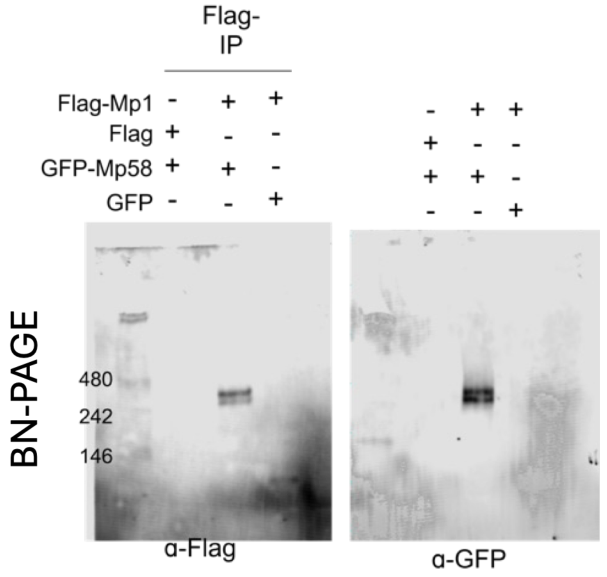


#### Rep2 (Fig. 3F)

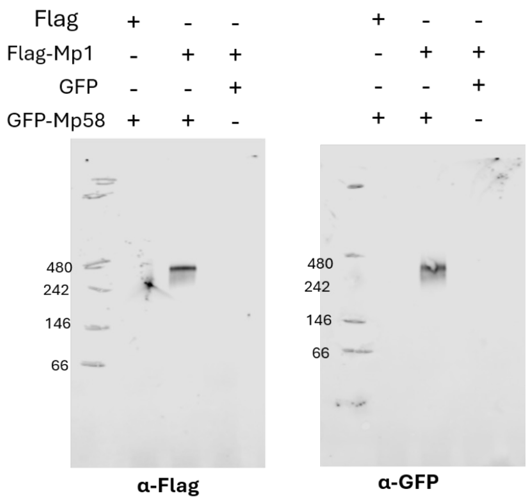

#### FLAG-Mp58$^{W66/F67E}$ x GFP-Mp1 co-IP

##### Rep1

##### Rep2 (Fig.6C)

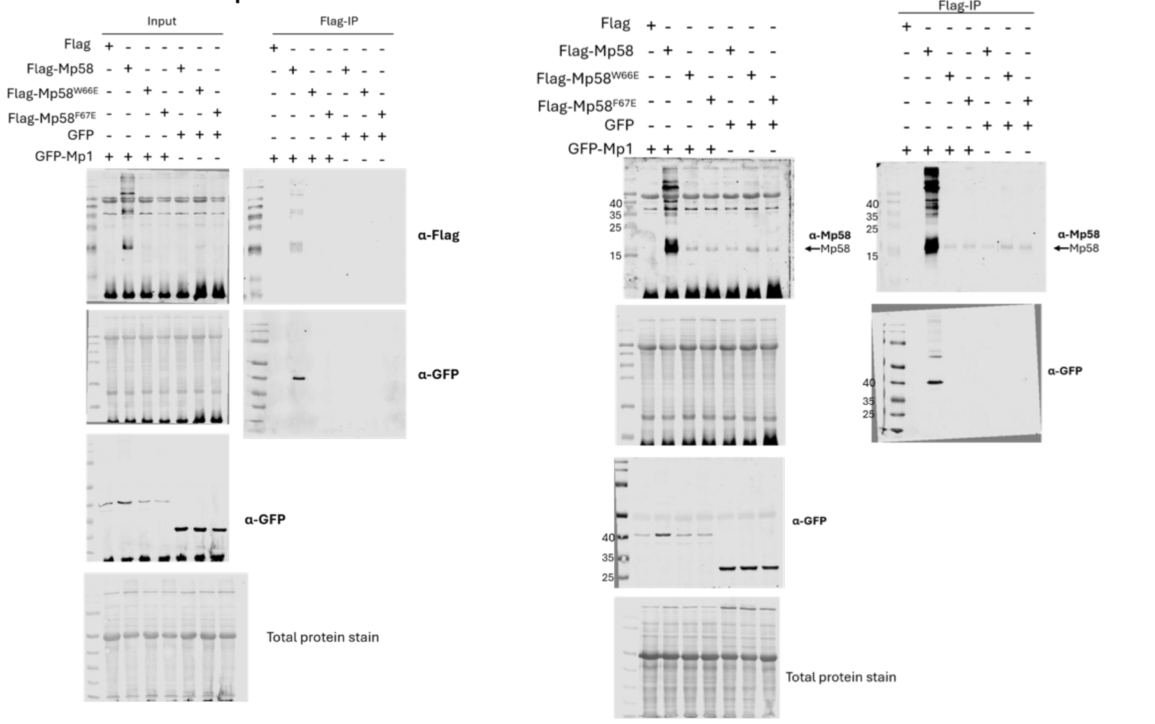


#### FLAG-Mp58$^{W66/F67A}$ x GFP-Mp1 co-IP (SDS-PAGE)

##### Rep1

##### Rep2 (Fig.6D)

##### Rep3

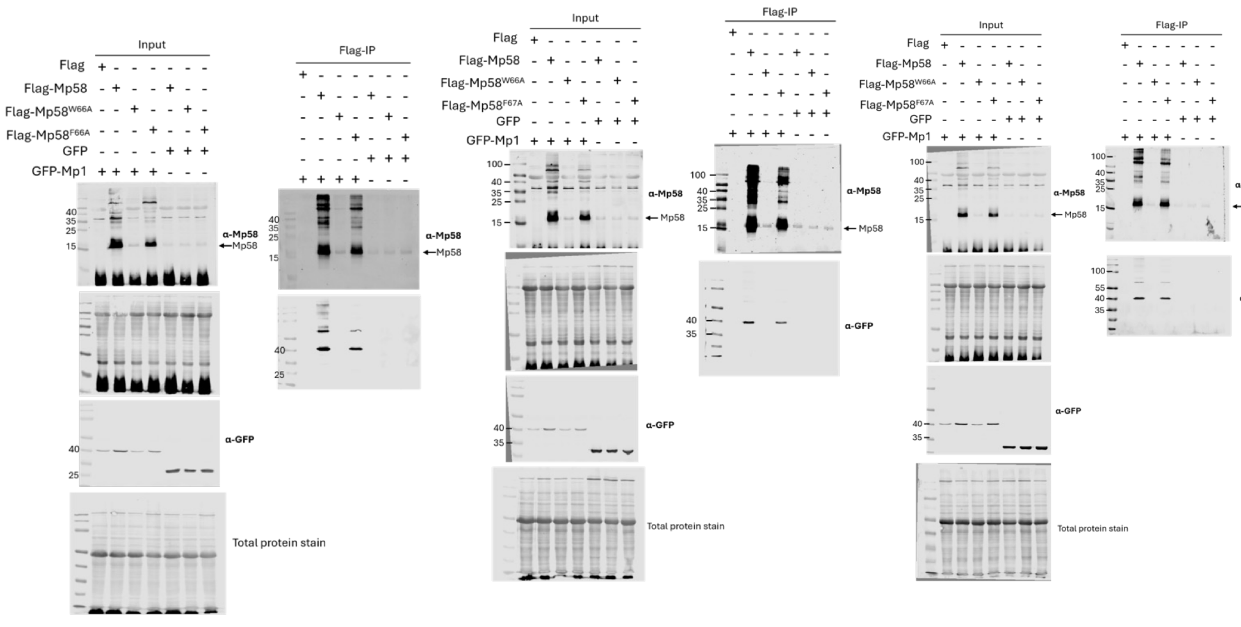

### FLAG-Mp58$^{W66/F67A}$ x GFP-Mp1 co-IP (BN-PAGE)

#### Rep1 (Fig.6E)

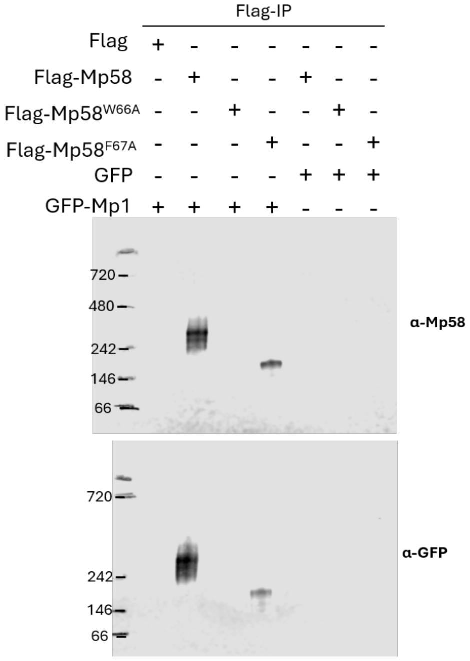


#### Rep2

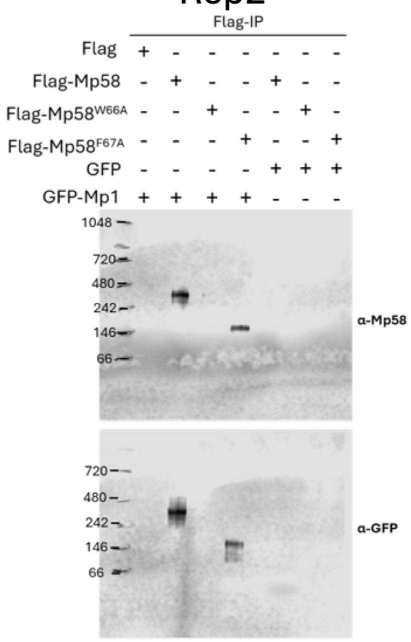


#### Fig S3A

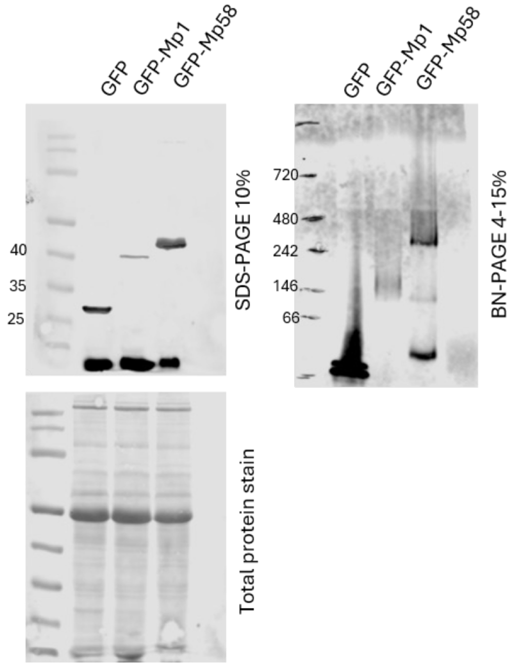


#### Fig S3B
#### (lanes 5&6 used in figure)

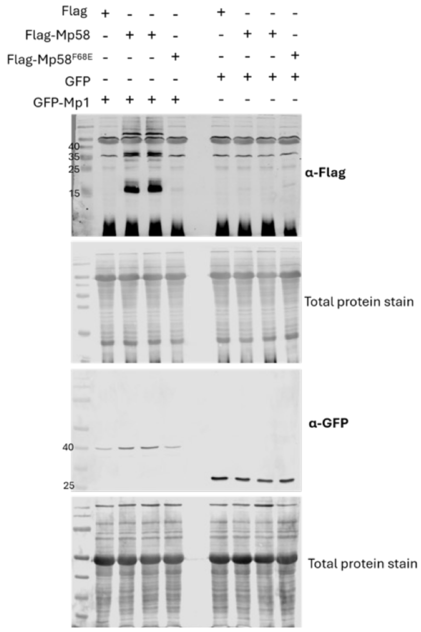
